# DNA damage signaling regulates cohesin stabilization and promotes meiotic chromosome axis morphogenesis

**DOI:** 10.1101/2021.08.28.458025

**Authors:** Zhouliang Yu, Abby F. Dernburg

## Abstract

A hallmark of meiosis is the reorganization of chromosomes as linear arrays of chromatin loops around a chromosome axis comprised of cohesins and regulatory proteins. Defective axis morphogenesis impairs homolog pairing, synapsis, and recombination. We find that axis assembly in *C. elegans* is promoted by DNA Damage Response (DDR) signaling activated at meiotic entry. Central to this regulation is downregulation of the cohesin release factor WAPL-1 by the DDR transducer kinase ATM-1, which is activated by the meiotic kinase CHK-2. Additional cohesin regulators, including ECO-1 and PDS-5, also contribute to stabilizing axis-associated cohesins. We find that downregulation of WAPL by ATM also promotes cohesin enrichment at DNA damage foci in cultured mammalian cells. Our findings reveal that the DDR and Wapl play conserved roles in cohesin regulation in meiotic prophase and proliferating cells.

**One-Sentence Summary:** DNA Damage Response kinase ATM phosphorylates WAPL to promote meiotic chromosome axis assembly and DNA repair

## Introduction

Sexually reproducing organisms produce haploid gametes via the specialized cell division process of meiosis. During meiotic prophase, homologous chromosomes pair and synapse to enable recombination and crossover formation, which underlie both reductional chromosome segregation and genetic variation (*1–3*). In early meiosis, replicated chromosomes become highly elongated as cohesins reorganize to form a linear chromosome “axis” (*4*). Axis morphogenesis is a prerequisite for the induction of meiotic double-strand breaks (DSBs), homolog pairing, synapsis, and DSB repair (*5–11*). Meiotic cohesins also recruit additional axis proteins that monitor synapsis and recombination and regulate cell cycle progression (*5, 7, 12–14*).

The remodeling of meiotic chromosomes to form an axis-loop structure is thought to be driven at least in part through the expression of meiosis-specific cohesin subunits. Cohesins are multisubunit protein complexes that form chromatin loops through their ATPase and DNA-binding activities. They are the most ancient and primary determinants of large-scale chromosome architecture (*15, 16*), in addition to maintaining connections between sister chromatids from the time of DNA replication to cell division. Establishment, maintenance, and regulated release of sister chromatid cohesion is essential for faithful chromosome segregation during mitosis and meiosis (*5, 6, 17*).

The core cohesin complex contains four components: a heterodimer of two large ATPases, the Structural Maintenance of Chromosomes (SMC) proteins Smc1 and Smc3; Sister Chromatid Cohesion 3/Stromal Antigen (Scc3/SA/Stag); and an α-kleisin protein. Kleisins are cleaved by a protease, separase, to enable chromosome segregation in mitosis and meiosis. All organisms studied to date express one or more meiosis-specific kleisins, and some also have meiosis-specific SMC and/or Scc3/Stag isoforms (*5, 6, 18*). However, it is largely unknown how the activities of meiotic cohesins differ from those in other cells, except that the Rec8 kleisin is protected from cleavage by separase to keep sister chromatids together during the first meiotic division.

Less attention has been paid to the potential meiotic roles of cohesin regulatory factors, which are thought to modulate the loading, unloading, and dynamics of cohesins on chromatin. These include the “loading” complex Scc2/Scc4, the acetyltransferase Eco1/Ctf7, the “cohesin release” protein Wapl, and Pds5, which plays essential but poorly understood roles in mitosis and meiosis. These factors are thought to modulate the ATPase activity of cohesin and the persistence of the complex on chromatin.

Here we investigate the mechanism of chromosome remodeling during early meiotic prophase in *C. elegans*. This nematode expresses two different meiotic kleisins: REC-8 and COH-3/4. COH-3 and COH-4 are closely related paralogs with overlapping roles and are thus regarded as a single type of kleisin. During most of meiotic prophase, cohesin complexes containing REC-8 and COH-3/4 localize along the length of chromosome axes. Both types are essential for homolog pairing and synapsis: COH-3/4 are more abundant and are thought to form an essential scaffold for synapsis, while REC-8 may fuse the axes of sister chromatids to prevent inter-sister recombination and synapsis (*9, 13, 19, 20*). Following crossover designation at mid-pachytene, the two types of cohesins become enriched on reciprocal “arms” of the bivalent to mediate reductional segregation (*9*). No other meiosis-specific cohesin proteins have been identified in *C. elegans*.

We show that WAPL-1 (Wapl) is downregulated by ATM-1 (Ataxia telangiectasia mutated, ATM) at meiotic entry, which is important to stabilize COH-3/4 along chromosome axes. Surprisingly, we find that ATM-1 is activated at meiotic entry by the CHK-2 kinase, and that CHK-2 also orchestrates a regulatory pathway that preferentially stabilizes REC-8. We extend our observations to show that inhibition of Wapl by ATM promotes cohesin enrichment at sites of DNA damage in proliferating human cells.

## Results

### WAPL-1 is downregulated at meiotic entry

Wapl is a widely conserved cohesin regulator that was identified in screens for radiation-sensitivity in budding yeast and mitotic defects in *Drosophila*. Its best-known role is to promote the release of cohesin along chromosome arms during mitotic prophase; Wapl does not cleave cohesin but stimulates its dissociation from chromatin. Intriguingly, depletion of Wapl from mammalian cells results in the formation of “vermicelli”, the clustering of cohesin to form axial structures that resemble meiotic chromosome axes, during interphase (*21*).

Immunolocalization of *C. elegans* WAPL-1 (Wapl) reveals diffuse nuclear localization in most tissues, including the germline. Interestingly, its abundance drops abruptly at meiotic entry (*22,23*). This reduction was more pronounced in dissected, immunostained gonads than in intact animals expressing GFP::WAPL-1, suggesting that WAPL-1 dissociates from chromosomes rather than being degraded in meiotic cells (*22*).

The formation of “vermicelli” upon WAPL depletion in mammals suggested that meiotic downregulation of WAPL-1 might contribute to axis assembly, perhaps through cohesin stabilization. WAPL-1 reaccumulates in oocyte nuclei during later stages of meiotic prophase and contributes to removing COH-3/4 cohesin from the axes as chromosomes condense. However, this role is dispensable for meiosis; *wapl-1* null mutants are viable and fertile, with normal meiotic segregation, as indicated by a normal frequency of male progeny, which arise through X chromosome nondisjunction. They do show hallmarks of reduced mitotic fidelity, including some embryonic and larval lethality and low-penetrance egg-laying and locomotion defects (*22, 23*).

We found that downregulation of WAPL-1 at meiotic entry requires the essential meiotic kinase CHK-2 (Fig. 1A). Either *chk-2* loss-of-function mutants or auxin-induced depletion of CHK-2 resulted in persistence of

**Fig. 1.**
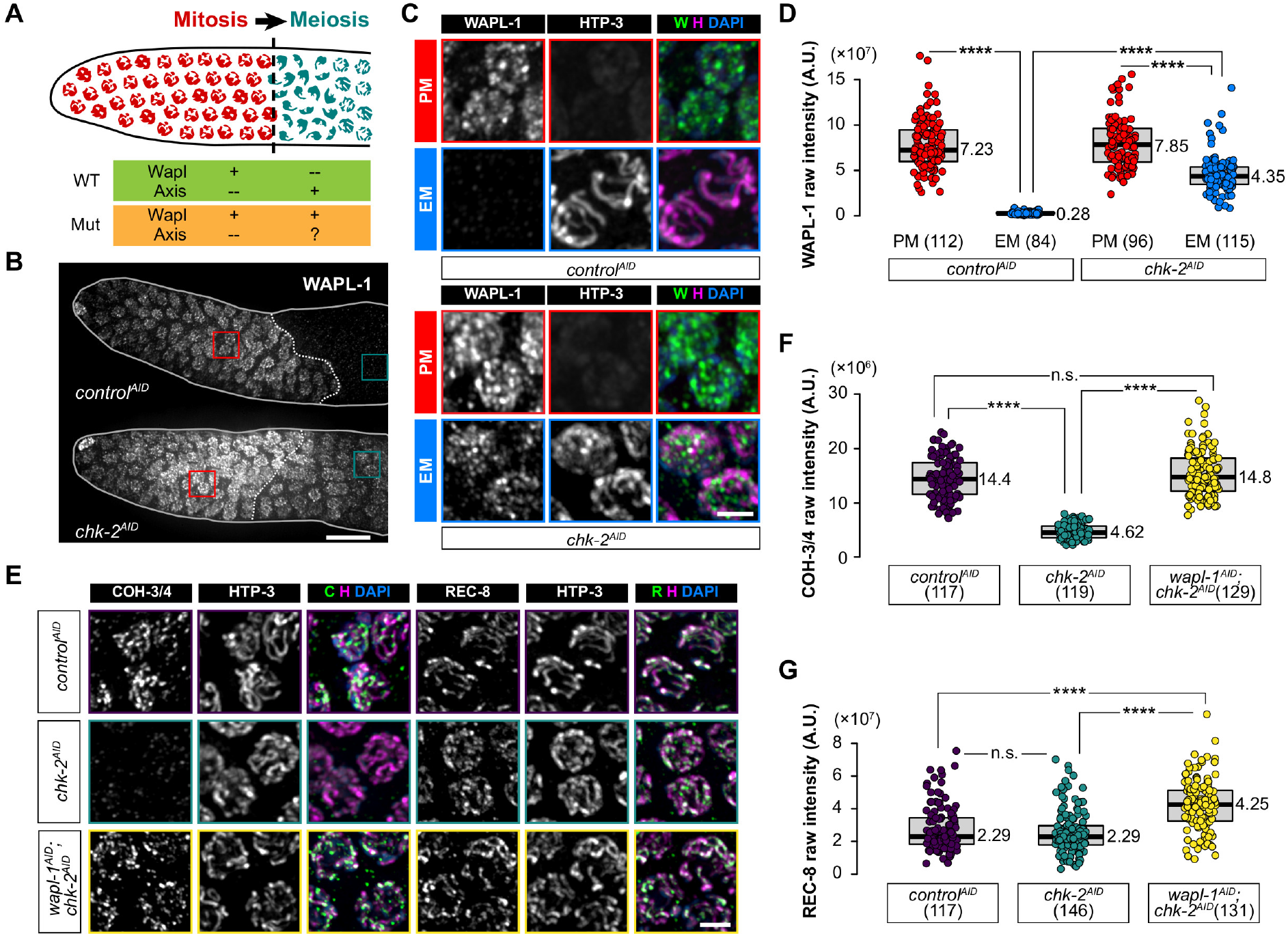
WAPL-1 is downregulated at meiotic entry. **(A)** Diagram of the distal tip of a *C. elegans* gonad containing proliferating germline (PM) stem cells (mitosis, in red) and early meiotic (EM) nuclei (leptotene/zygotene/early pachytene, in blue). The dashed line indicates the boundary between premeiotic and meiotic cells. **(B)** WAPL-1 immunostaining in the distal tip of gonads from auxin-treated animals. Red and blue rectangles outline regions containing premeiotic and early meiotic nuclei, respectively, which are enlarged in **(C).** Scale bar, 10 μM. Meiotic nuclei display chromosome marked by the HORMA domain protein HTP-3 (magenta). Scale bar, 2 μM. **(D)** Quantification of WAPL-1 immunostaining in (B). **(E)** COH-3/4 and REC-8 immunolocalization (in green) in early pachytene nuclei. Scale bar, 2 μM. **(F)** and **(G)** Quantification of the intensity of COH-3/4 and REC-8 immunostaining in (E) See “Data Presentation” for details.

WAPL-1 in early meiotic nuclei (Fig. 1B and 1D). Loss of CHK-2 does not abolish axis assembly or recruitment of the HORMA domain protein HTP-3 (Fig. 1C). However, the abundance of COH-3/4 kleisins was greatly reduced in the absence of CHK-2, consistent with prior evidence that WAPL-1 preferentially releases COH-3/4-containing cohesins from meiotic chromosomes (Fig. 1E and 1F) (*23*). Co-depletion of WAPL-1 and CHK-2 fully restored COH-3/4 to wildtype levels (Fig. 1E and 1F). Interestingly, we also observed a significant increase in REC-8 intensity following this co-depletion (Fig. 1E and 1G), similar to observations in *wapl-1* mutants (*24*).

WAPL-1 contains four consensus phosphorylation sites for CHK-2 (R-[x]-[x]-S/T) Surprisingly, mutation of all four sites to nonphosphorylatable residues (*wapl-1^4SA^*) did not affect the localization of WAPL-1 in proliferating or meiotic nuclei (Figures S1A-S1D). Thus, we considered the possibility that the regulation of WAPL-1 by CHK-2 might be indirect. We tested whether WAPL-1 downregulation requires the formation of meiotic DSBs, synapsis, and/or homologous pairing, three central meiotic events that depend on CHK-2 activity in early prophase (*25–27*). We depleted SPO-11, which catalyzes meiotic DSB formation, and SYP-3, an essential SC component, but neither affected the reduction of WAPL-1 in early prophase (Fig. S1E-G) (*28, 29*). WAPL-1 also showed normal reduction in a *plk-2; plk-1* double deplete, which has greatly reduced homologous pairing (data not shown) (*30, 31*).

Previous studies have shown that the DNA Damage Response (DDR) transducer kinase ATM mediates a genome-wide increase of cohesin binding in response to ionizing radiation (IR)-induced DNA damage in mammals (*32*). DDR signaling also governs localized cohesin loading at DNA damage loci (*33–36*). Additionally, Wapl was identified as an ATM/ATR target in *Arabidopsis* (*37*). We found that depletion of ATM-1 (ATM) in the germline completely abolished WAPL-1 downregulation, whereas depletion of ATL-1 (ATR) had no effect (Fig. 2B-2D). Alignment of WAPL-1 homologs also revealed a potentially small but conserved cluster of S/T-Q residues (Fig. 2A). Such S/T-Q cluster domains (SCDs) are found in many ATM/ATR substrates. We replaced the two serines in the WAPL-1 SCD with nonphosphorylatable (*wapl-1^2A^*) or phosphomimetic (*wapl-1^2D^*) amino acids (Fig. 2A). WAPL-1^2A^ showed defective WAPL-1 downregulation (Fig. 2E), despite normal levels of CHK-2 (Fig. S2A-S2C) and ATM-1 (Fig. S2D-S2F). By contrast, a phosphomimetic WAPL-1^2D^ showed substantial reduction of WAPL-1 both before and after meiotic entry (Fig. 2E). These results support the idea that ATM-1 downregulates WAPL-1 by phosphorylating its N-terminal SCD. Localization of WAPL-1^2D^ was not restored by depletion of CHK-2 or ATM-1 (Fig. 2E-2G).

**Fig. 2.**
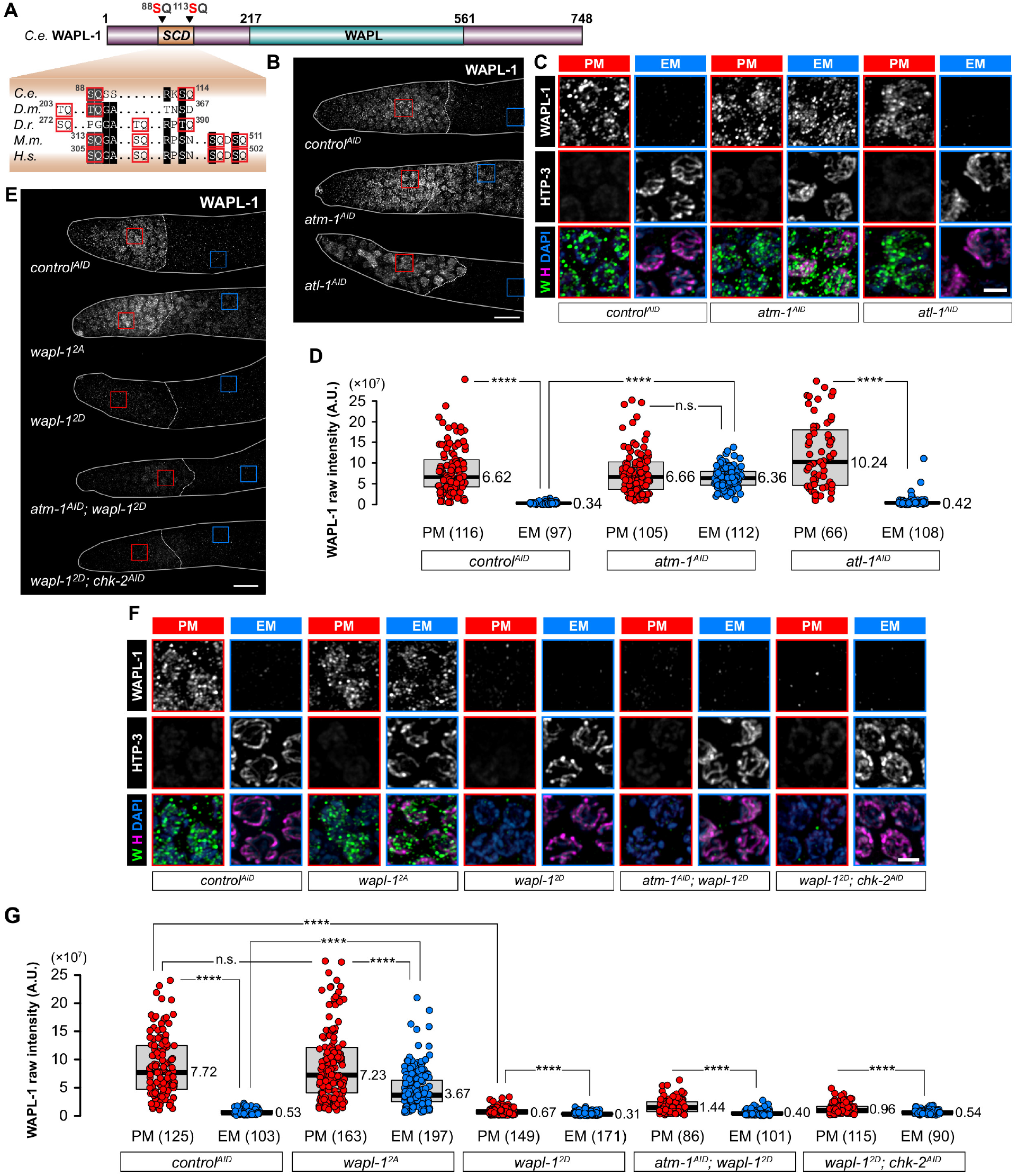
WAPL-1 suppression depends on ATM-1 and a SCD. **(A)** Diagram illustrating the domain architecture of *C. elegans* WAPL-1. The N-terminal SCD identified in this study and the two putative phosphorylation sites are shown. The partial alignment of amino acid sequences of corresponding regions of Wapl orthologs show conservation of this small SCD across species (*C.e., Caenorhabditis elegans; D.m., Drosophila melanogaster; D.r., Danio rerio; M.m., Mus musculus; H.s., Homo sapiens*). SQs and TQs are highlighted and outlined. **(B)** and **(E)** WAPL-1 immunostaining in the distal tip of gonads. Scale bar, 10 μM. **(C)** and **(F)** Enlarged images showing WAPL-1 immunostaining (in green) in premeiotic nuclei and early meiotic nuclei from (B) and (E). HTP-3 is recruited to axes at meiotic entry. Scale bar, 2 μM. **(D)** and **(G)** Quantification of the intensity of WAPL-1 immunostaining in (B) and (E).

### CHK-2 positively regulates ATM-1 activity

In the canonical DNA damage response pathway in mammalian cells, Chk2 is an essential downstream mediator of ATM activity (*38, 39*). However, previous studies have shown that *C. elegans* CHK-2 is dispensable for checkpoint activation in response to hydroxyurea and ionizing radiation in embryos and the adult germline, and is only essential for meiosis (*25, 26*). Like other meiosis-specific Chk2 orthologs, *C. elegans* CHK-2 lacks the N-terminal SCD that normally mediates activation by ATM (Fig. 3A). Additionally, CHK-2-dependent phosphorylation of the nuclear envelope protein SUN-1 was observed in the absence of ATM-1 and ATL-1 (*40*). We further found that pairing center proteins HIM-8 and ZIM-1, −2, and −3, which are direct targets of CHK-2, were still phosphorylated when we depleted ATM-1, ATL-1, or both proteins (Fig. 3B and 3C), confirming that CHK-2 activity is independent of ATM-1 and ATL-1.

**Fig. 3.**
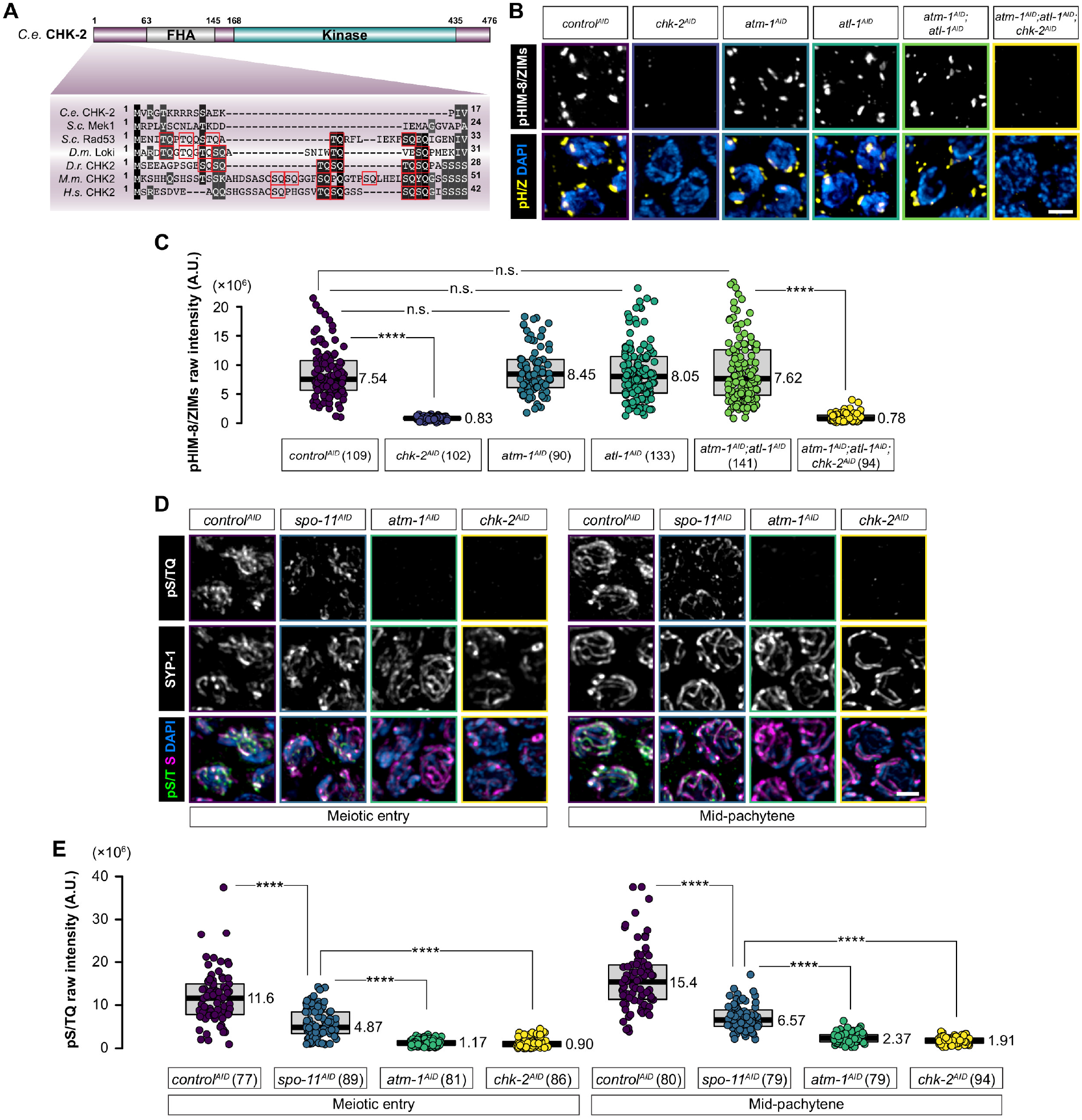
CHK-2 activates ATM-1 to suppress WAPL-1 during early meiotic prophase. **(A)** Domain architecture of *C. elegans* CHK-2. Partial alignment of the N-termini of several Rad53/CHK2 orthologs (*C.e., Caenorhabditis elegans; S.c., Saccharomyces cerevisiae; D.m., Drosophila melanogaster; D.r., Danio rerio; M.m., Mus musculus; H.s., Homo sapiens*) is shown below the schematic, with all S/TQs highlighted. **(B)** Immunofluorescence using a phospho-specific antibody that recognizes CHK-2 target motifs in HIM-8 and ZIM-1, −2, and −3. Scale bar, 2 μM. **(C)** Quantification of pHIM-8/ZIM immunostaining in (B). **(D)** pS/TQ immunostaining (in green) of meiotic entry nuclei (left) or mid-pachytene nuclei (right) under indicated conditions. SYP-1 immunostaining (in magenta) shows the SC. Scale bar, 2 μM. **(E)** Quantification of the intensity of pS/TQ immunostaining in (D).

Evidence that CHK-2 activity is independent of ATM-1 and WAPL-1 persists on chromatin in the absence of either CHK-2 or ATM raised the possibility that ATM-1 activity might depend on CHK-2. We tested this, first using a phospho-specific antibody that recognizes ATM substrates (pS/TQ) (*41*), and found that the signal in early meiotic nuclei was greatly reduced following depletion of either CHK-2 or ATM-1 (Fig. 3D).

Formation of meiotic DSBs in *C. elegans* requires CHK-2 (*26, 42, 43*). We thus tested whether ATM-1 activity requires DSBs by depleting SPO-11, and found that this partially reduced pS/TQ immunofluorescence, but less so than depletion of ATM-1 (Fig. 3D and 3E). This is consistent with our evidence that SPO-11 depletion did not affect WAPL-1 downregulation at meiotic entry (Fig. S1A-S1C). Together, these results reveal that CHK-2 promotes basal levels of ATM-1 activity that mediate WAPL-1 downregulation even in the absence of meiotic DSBs.

### A conserved CHK-2 consensus site in the FAT domain is essential for ATM-1 activity

Our results suggested that CHK-2 might directly regulate ATM-1 activity independently of meiotic DSB formation. Previous studies have shown that high concentrations of Chk2 can lead to its self-activation *in vitro* and phosphorylation of H2AX and other S/T-Q sites *in vivo* (*44, 45*), suggesting that Chk2 may promote ATM/ATR activity under certain conditions. We aligned the amino acid sequences of ATM family proteins and found a conserved CHK-2 consensus phosphorylation motif (R-[x]-[x]-S/T) within their FAT domains, which are critical for ATM/ATR activation (Fig. 4A) (*46–48*). A mutation in this motif was also identified in a leukemia patient with ATM deficiency (*49*).

**Fig. 4.**
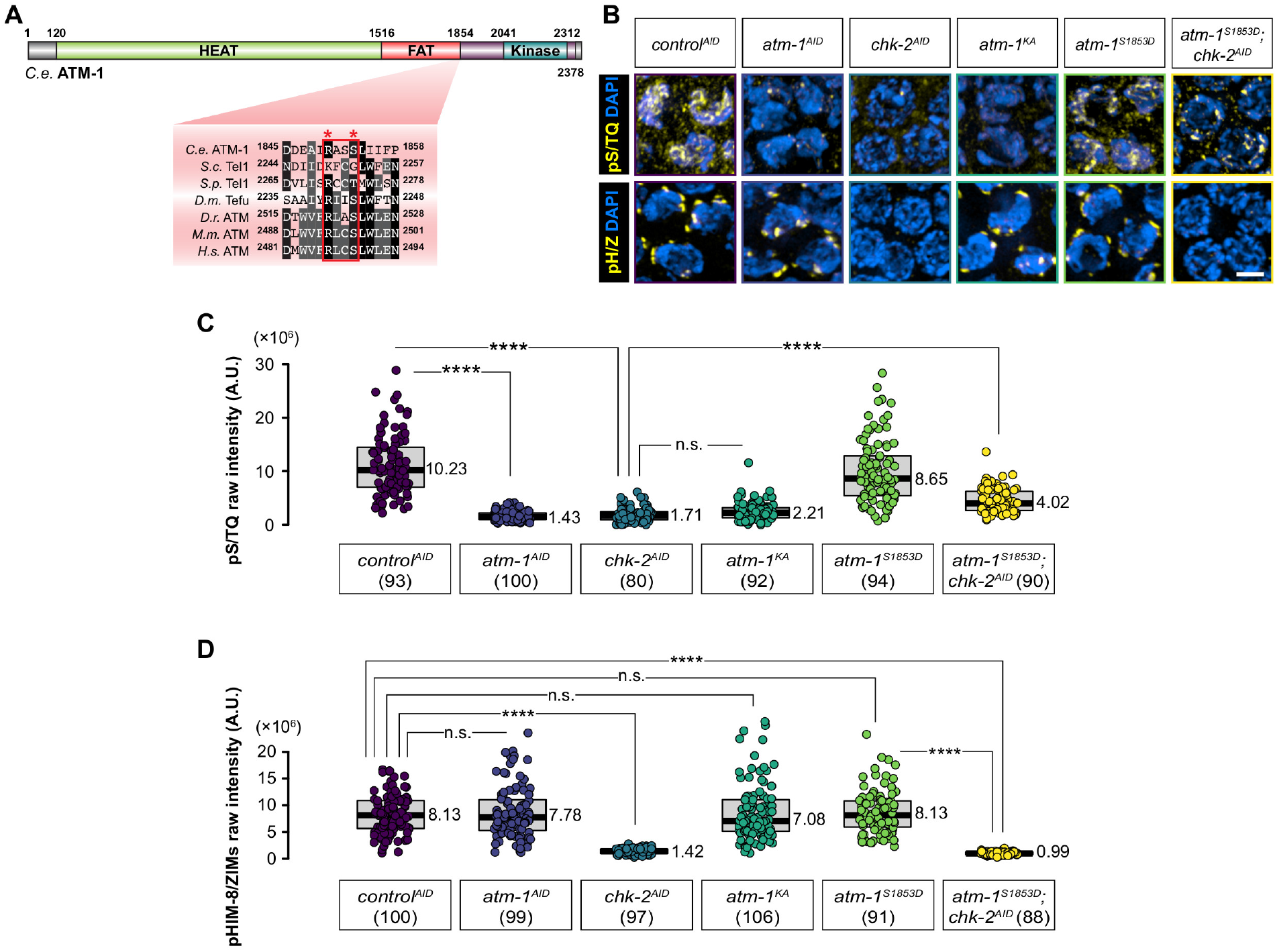
A CHK-2 consensus motif in the FAT domain of ATM-1 mediates ATM-1 activity. **(A)** Domain architecture of *C. elegans* ATM-1. The alignment of the C-terminal end of FAT domains from several ATM orthologs (*C.e., Caenorhabditis elegans; S.c., Saccharomyces cerevisiae; S.p., Schizosaccharomyces pombe; D.m., Drosophila melanogaster; D.r., Daniorerio; M.m., Mus musculus; H.s., Homo sapiens*) is shown below the schematic, with the Rad53/CHK2 consensus motif outlined. The conserved arginine and phospho-serine/threonine site of the consensus motif are indicated with asterisks. **(B)** pS/TQ immunostaining is used as a proxy for ATM-1/ATR-1 activity, while phosphorylation of conserved motifs on HIM-8 and the ZIM proteins is indicative of CHK-2 activity. Scale bar, 2 μM. **(C)** and **(D)** Quantification of of pS/TQ immunofluorescence intensity (C) and pHIM-8/ZIM intensity (D) (see Data Presentation for details).

Mutation of this putative phosphosite (*atm-1^KA^*) greatly reduced pS/T-Q immunofluorescence in meiotic nuclei (Fig. 4A and 4B), while the corresponding phosphomimetic mutation (*atm-1^S1853D^*) resulted in normal ATM-1 activity (Fig. 4A and 4B). Additionally, *atm-1^S1853D^* showed high ATM-1 activity even in the absence of CHK-2, supporting the conclusion that CHK-2 activates ATM-1 through this site (Fig. 4B-4D).

These findings indicate that the persistence of WAPL-1 in meiotic nuclei lacking CHK-2 is a consequence of a failure to activate ATM-1. To further test this idea,we examined WAPL-1 immunostaining in *atm-1^KA^* and *atm-1^S1853D^*, alleles that showed defective and normal kinase activity of ATM-1, respectively. CHK-2 activity in these alleles was normal (Fig. 4B-4D). Animals with only the nonphosphorylatable *atm-1^KA^* allele showed aberrant WAPL-1 staining, similar to that seen following depletion of CHK-2 or ATM-1 (Fig. 5A-5C). By contrast, the phosphomimetic *atm-1^S1853D^* allele resulted in normal downregulation of WAPL-1, even when CHK-2 was depleted (Fig. 5A-5C). We tested whether *atm-1^S1853D^* could bypass the requirement for CHK-2 in DSB induction, and found that RAD-51 foci were absent, indicating that CHK-2-dependent ATM activation is dispensable for the formation of meiotic DSBs (Fig. S4A-S4B).

**Fig. 5.**
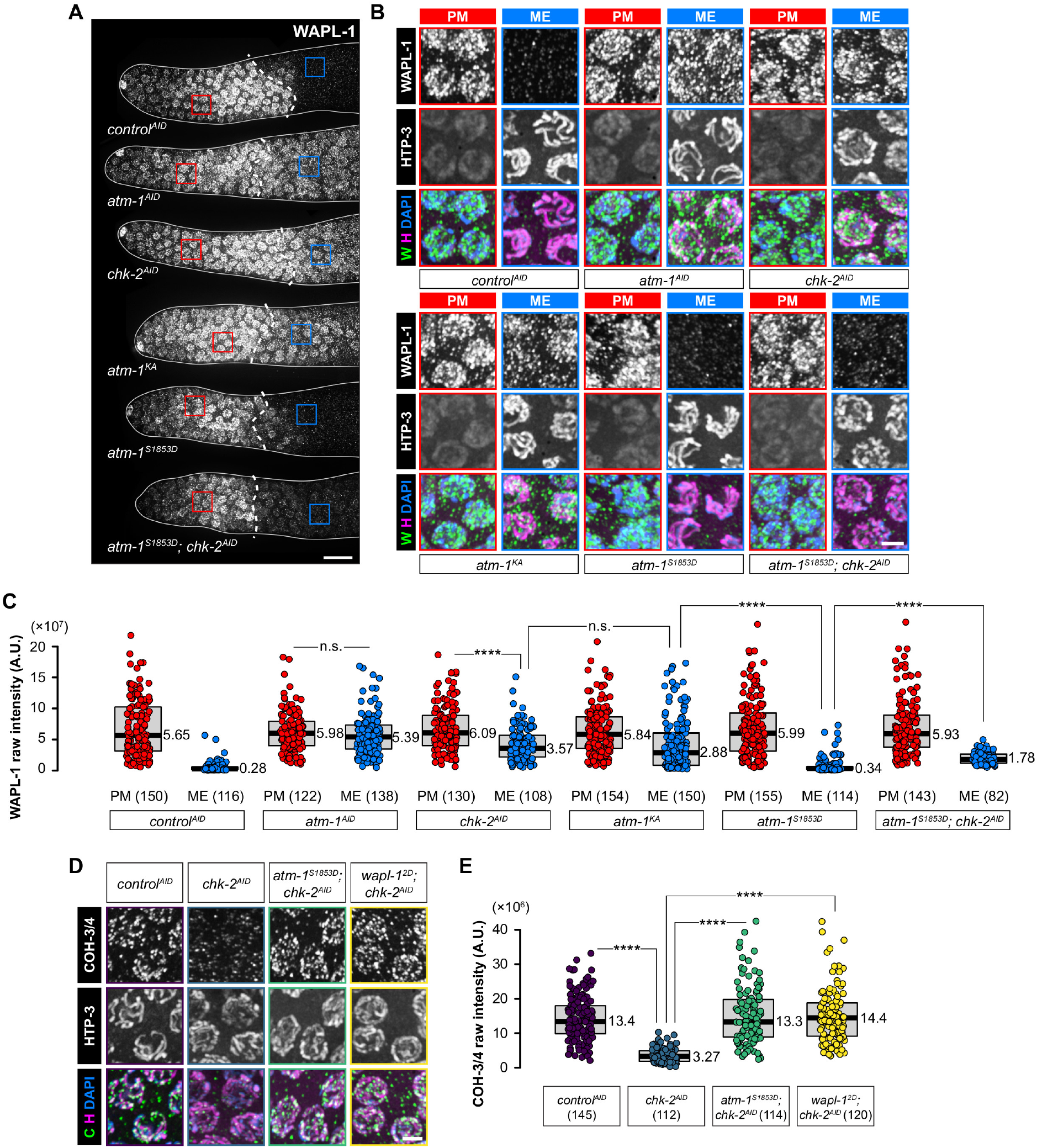
CHK-2 suppresses WAPL-1 by activating ATM-1. **(A)** WAPL-1 immunostaining in the distal tip of gonads. Scale bar, 10 μM. **(B)** Enlarged images of the regions indicated in (A). HTP-3 localizes to axes starting at meiotic entry. Scale bar, 2 μM. **(C)** Quantification of the intensity of WAPL-1 immunostaining in (A). **(D)** COH-3/4 immunostaining (in green) of early pachytene nuclei. Scale bar, 2 μM. **(E)** Quantification of the intensity of COH-3/4 immunostaining in (D).

Together, our results indicated that CHK-2 activates ATM even in the absence of DSBs (Fig. 3D and 3E), and that this CHK-2-dependent ATM-1 activity down-regulates WAPL-1 to stabilize cohesins. Phosphomimetic mutations in ATM-1 or WAPL-1 were sufficient for robust axis localization of COH-3/4 in the absence of CHK-2 (Fig. 5D and 5E).

### CHK-2 also promotes axis assembly through cohesin acetylation

The evidence above indicates that CHK-2 regulates COH-3/4 localization through ATM-1 and WAPL-1. However, COH-3/4 was more severely reduced along the axis in the absence of CHK-2 than in the absence of ATM-1 or expression of a nonphosphorylatable allele of WAPL-1 (Fig. S5A), suggesting that CHK-2 may also promote other activities that stabilize COH-3/4. Our evidence that COH-3/4 localization is fully restored by WAPL-1 depletion in *chk-2* mutants indicated that CHK-2 may promote a pathway that antagonizes WAPL-1 activity. Factors known to antagonize Wapl include Sororin in vertebrates and *Drosophila* (*50–52*), and the more broadly conserved acetyltransferase Eco1/ESCO1/ECO-1 (*53–56*), which can render cohesins insensitive to mobilization by Wapl.

We found that depletion of ECO-1 alone had no effect on axial COH-3/4 localization (Fig. 6A-6C). However, co-depletion of ECO-1 and ATM-1 or depletion of ECO-1 in *wapl-1^2A^* showed synergistic effects on COH-3/4 localization (Fig. S5A-S5C), nearly recapitulating the effects of CHK-2 depletion (Fig. 6A-6C). These results suggested that ECO-1 can antagonize WAPL-1 in early meiosis, but that WAPL-1 downregulation normally makes this unnecessary.

**Fig. 6.**
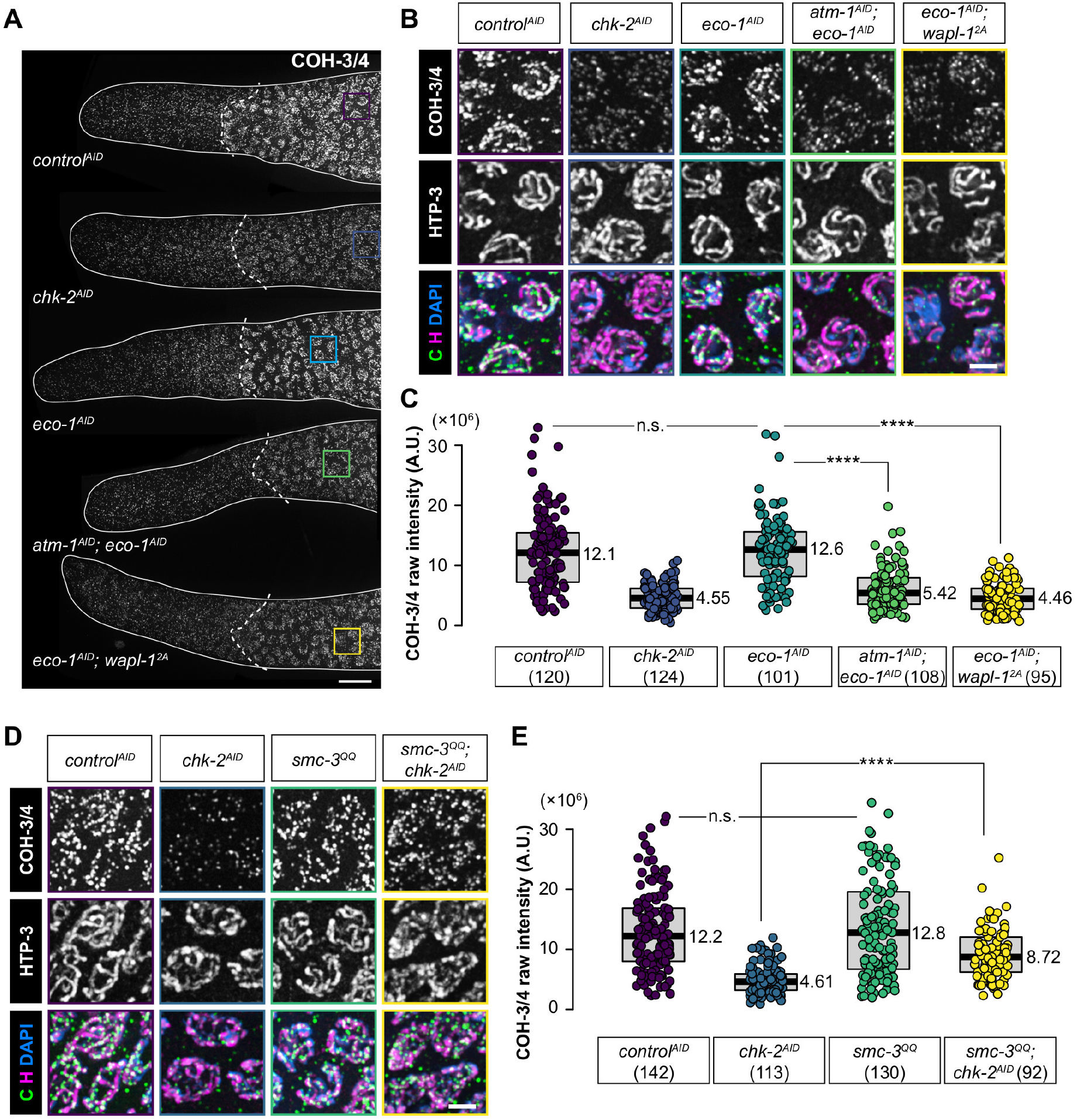
ECO-1-dependent cohesin acetylation contributes to stabilization of axial cohesin against WAPL-1. **(A)** COH-3/4 immunostaining in the distal tip of gonads. Scale bar, 10 μM. **(B)** Enlargements of the regions indicated in (A). Bright HTP-3 staining along axes is detected starting at meiotic entry. Scale bar, 2 μM. **(D)** COH-3/4 immunostaining (in green) of meiotic entry nuclei. Scale bar, 2 μM. **(C)** and **(E)** Quantification of the intensity of COH-3/4 immunostaining in (B) and (D), respectively.

Previous studies have shown that Eco1/Eso1/ESCO1/2 antagonizes Wapl-dependent cohesin release by acetylating cohesin subunits (*57, 58*), including two conserved lysine sites on the ATPase head of Smc3 (*59–62*). We mutated the corresponding lysines in *C. elegans* SMC-3 to glutamine to mimic acetylation (K106Q/K107Q). Immunostaining using a commercial antibody against SMC-3 confirmed the expression and axial localization of SMC-3^QQ^ in meiotic cells. (data not shown). Axial COH-3/4 localization and axis morphogenesis appeared normal in early meiosis in *smc-3^QQ^* mutants (Fig. 6D). This mutation partially restored COH-3/4 localization in the absence of CHK-2 (Fig. 6D and 6E). These results suggest that CHK-2 promotes cohesin stabilization through at least two mechanisms, including downregulation of WAPL-1 and acetylation by ECO-1.

### PDS-5 protects REC-8 from WAPL-mediated release

We primarily used COH-3/4 localization as a marker to investigate the regulation of axis assembly by WAPL-1, since prior work and our observations showed more subtle effects of *wapl-1* mutations on REC-8 localization (Fig. 1E-1G) (*23*). However, Wapl can promote release of Rec8 cohesin in yeast, plant and human meiocytes (*63–66*). Therefore, we wondered why *C. elegans* REC-8 cohesin is more resistant to WAPL-1 activity than COH-3/4 during meiotic prophase.

The cohesin regulator Spo76/EVL-14/Pds5/PDS-5 is required to establish and maintain cohesion in mitosis and meiosis from yeast to human (*67–70*). Pds5 binds directly to kleisins, making it a plausible candidate for kleisin-specific activity during meiosis. Intriguingly, Pds5 can either recruit Wapl to release cohesin or prevent Wapl from accessing cohesin, depending on the context (*52, 56, 67, 69, 71*). Importantly, chromosome condensation defects seen in budding yeast *PDS5* mutants are rescued by loss of Rad61/Wpl1 (Wapl), suggesting a direct antagonism between Pds5 and Wapl (*72, 73*).

PDS-5 (EVL-14) is essential for gonad development and fertility in *C. elegans*, presumably due to its mitotic functions (*70*). The protein localizes to nuclei throughout the germline and is enriched along chromosome axes in meiotic nuclei (Fig. S6A and S6B). Its abundance on axes was not affected by loss of CHK-2 (Fig. S6A-S6C), despite the reduction in COH-3/4 (above), supporting the idea that PDS-5 preferentially associates with REC-8 cohesin.

Depletion of PDS-5 did not affect the abundance of REC-8 in premeiotic nuclei, but in early meiosis REC-8 was markedly reduced along axes (Fig. 7A, 7B, and 7E). This is similar to findings in fission yeast but contrasts with findings from budding yeast and mammals, where loss of Pds5 has little effect on Rec8 binding to chromosomes (*74–78*). By contrast, the localization of COH-3/4 cohesin was unaltered by depletion of PDS-5, supporting the idea that PDS-5 preferentially stabilizes REC-8 (Fig. 7C, 7D, and 7F).

**Fig. 7.**
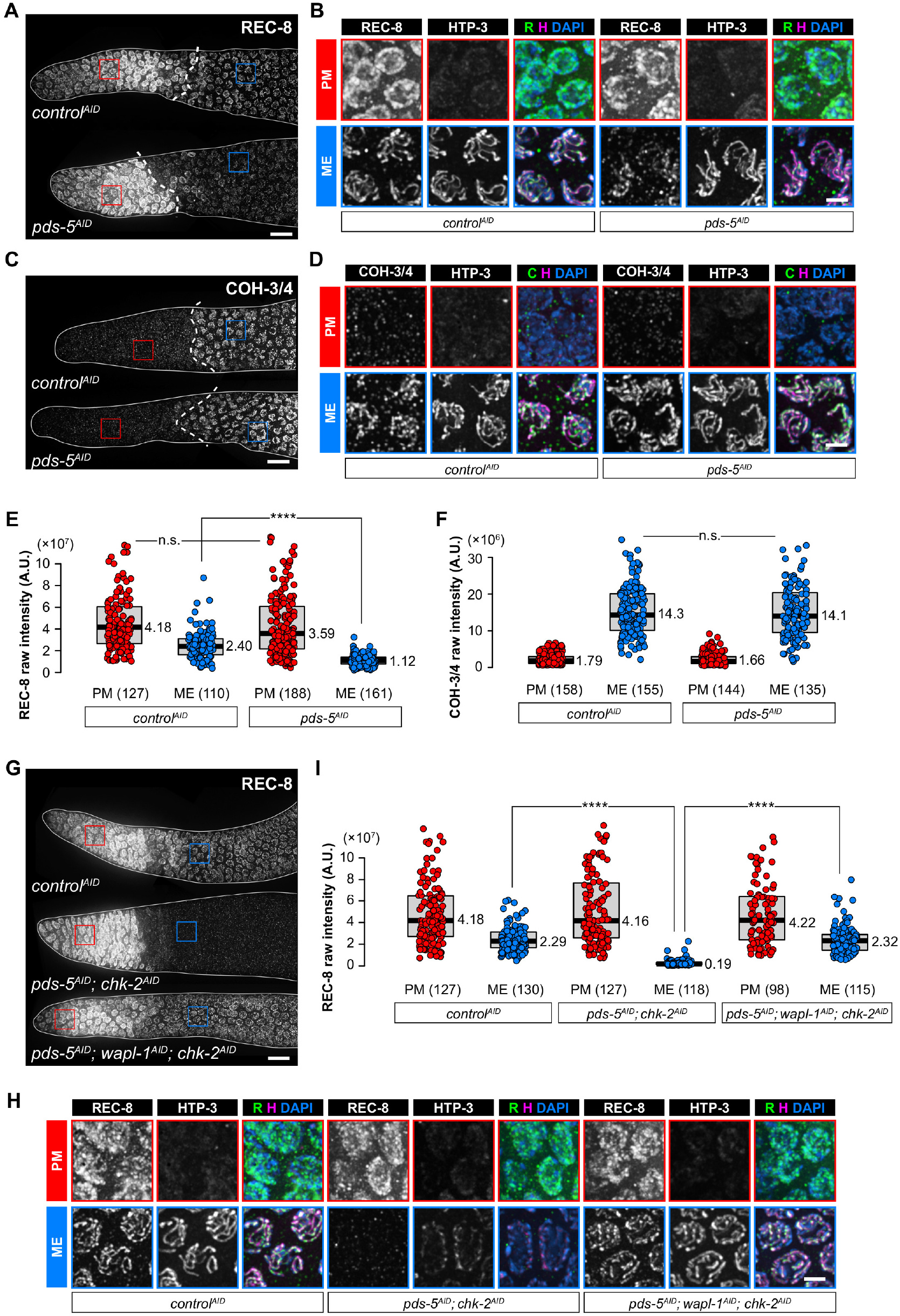
PDS-5 protects REC-8 cohesin from WAPL-1-dependent release in early meiosis. **(A) (C)** and **(G)** REC-8 (A and G) and COH-3/4 (C) immunofluorescence in the distal tip of gonads. Scale bar, 10 μM. **(B) (D)** and **(H)** Enlargements of the regions indicated in (A) (C) and (G), respectively. Scale bar, 2 μM. **(E) (F)** and **(I)** Quantification of the intensity of REC-8 (E and I) and COH-3/4 (F) immunostaining.

Codepletion of CHK-2 and PDS-5 resulted in dramatic reduction of REC-8 along chromosome axes (Fig. 7G), although premeiotic REC-8 was unaffected (Fig. 7G-7H). Importantly, REC-8 localization in the absence of PDS-5 and CHK-2 was fully rescued by codepletion of WAPL-1, indicating that PDS-5 protects REC-8 from release by WAPL-1, which becomes more apparent when WAPL-1 is not properly downregulated at meiotic entry (Fig. 7G-7I). The HORMA domain protein HTP-3 also failed to be recruited to axes in the absence of CHK-2 and PDS-5, consistent with evidence that HTP-3 recruitment requires cohesins but does not specifically require either REC-8 or COH-3/4 (Fig. 7G-7I) (*9*). Importantly, axial localization of COH-3/4, REC-8, and HTP-3 was restored by co-depletion of WAPL-1 (Fig. 7G-7I). Together these results demonstrate that downregulation of WAPL-1 and protection of REC-8 by PDS-5 are parallel pathways that contribute to axis assembly at meiotic entry.

### ATM-dependent WAPL downregulation in human cells promotes cohesin concentration at DNA damage foci

Our discovery of a conserved SCD in Wapl family proteins prompted us to explore whether ATM-dependent WAPL downregulation is important in contexts other than meiosis. Studies in yeast and human have shown that DDR transducer kinases ATM/ATR mediate cohesin recruitment at DNA damage loci to promote repair (*32, 36, 79*). ATM has also been found to promote the enrichment of CTCF, a cohesin binding partner, to DNA damage sites in human cells (*80*). Therefore, we investigated whether the establishment of the cohesin-enriched domain at DNA damage foci, where ATM is highly active, reflects ATM-mediated WAPL downregulation.

We treated HeLa cells with DNA damage-inducing agents, including the radiomimetic DNA-cleaving agent bleomycin (BLEO) (*81*) and the topoisomerase II poison etoposide (ETO) (*82*), both of which give rise to DSBs that activate ATM. Upon treatment with either bleomycin or etoposide, we observed the appearance of foci positive for both γH2A.X (S139-phosphorylated histone H2A.X) (Fig. 8A) (*83, 84*) and pS/TQ (Fig. 8B) (*41*). The mitotic kleisin Rad21 concentrated at many of these damage foci (Fig. 8A-8D).

**Fig. 8.**
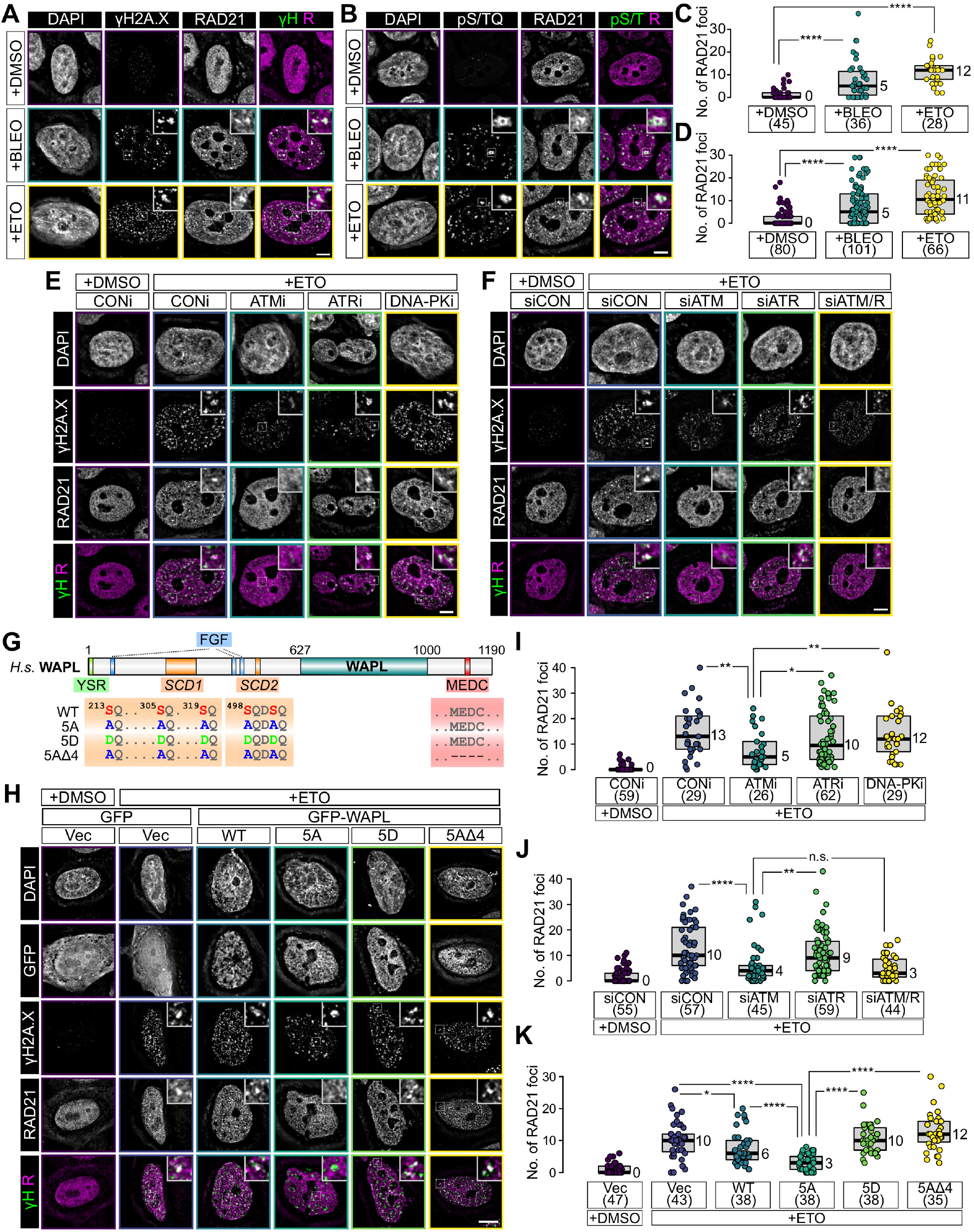
ATM-mediated WAPL downregulation regulates cohesin concentration at DNA damage foci. **(A)** and **(B)** RAD21 immunostaining in nuclei of HeLa cells that are treated with either bleomycin (BLEO) or etoposide (ETO). Scale bar, 10 μM. **(C)** and **(D)** Quantification of the number of RAD21 foci under conditions shown in (A) and (B). **(E)** RAD21 immunostaining in nuclei of HeLa cells treated with kinase inhibitors, followed by etoposide (ETO) to induce DNA damage. Chemical inhibitors specific for different kinases were KU55933 (ATMi), VE-821 (ATRi), and NU7441 (DNA-PKi). Scale bar, 10 μM. **(F)** RAD21 immunostaining in nuclei of HeLa cells depleted of ATM and/or ATR prior to etoposide (ETO)-induced DNA damage. Scale bar, 10 μM. **(G)** Domain architecture of human WAPL, indicating the positions of residues mutated in our transgenic constructs. **(H)** Immunofluorescence of HeLa cell nuclei expressing GFP or GFP-WAPL. Scale bar, 10 μM. **(I) (J)** and **(K)** Quantification of the number of RAD21 foci in (E) (F) and (H), as described under Image Analysis and Data Presentation.

We next induced damage in cells in the presence of specific inhibitors of ATM, ATR and DNA-PK (Fig. 8E), three paralogous DNA damage transducer kinases. Inhibition of ATM, but not ATR or DNA-PK resulted in markedly reduced γH2A.X upon etoposide treatment (Fig. 8E and S7A). Inhibition of ATM also largely eliminated damage-induced Rad21 foci (Fig. 8E and 8I).

To corroborate the specificity of the chemical inhibitors, we performed siRNA-mediated knockdown of ATM and ATR (Fig. 8F). We confirmed knockdown of ATM using a commercial antibody (Fig. S7C and S7D). Although we did not have a similar tool to monitor ATR abundance, we noted that nuclear volume was reduced by either ATR inhibition or ATR knockdown (Fig. S7B and S7G). Consistent with our observations with small-molecule inhibitors, ATM knockdown resulted in a significant decrease in nuclear γH2A.X intensity (Fig. 8F, S7F). Importantly, Rad21 enrichment was no longer detected at damage foci marked by either γH2A.X or pS/TQ (Fig. 8F, 8J and S7E). Together, these results indicated that the activity of ATM, but not ATR or DNA-PK, is required for cohesin enrichment at DNA damage foci in HeLa cells.

To test whether ATM-mediated WAPL downregulation contributes to the assembly of these cohesin-enriched domains, we designed nonphosphorylatable and phosphomimetic mutations at the most likely ATM phosphorylation sites in Wapl. Human Wapl has two potential SCDs (Fig. 8G). Importantly, neither SCDs overlaps with the FGF or YSR domains, which mediate the interaction between WAPL and PDS5 (*85, 86*). Overexpression of GFP-tagged wild-type WAPL or mutant proteins did not affect the appearance of γH2A.X following DNA damage (Fig. 8G-8H), indicating that ATM signaling is intact upon WAPL overexpression (Fig. 8H and S7H). However, overexpression of wild-type WAPL reduced nuclear RAD21 foci following damage (Fig. 8H and 8K). Overexpression of nonphosphorylatable WAPL (WAPL^5A^) blocked the formation of RAD21 foci even more effectively, but WAPL^5D^ lacked this activity (Fig. 8H, 8K). To test whether the reduction of RAD21 foci upon WAPL^5A^ overexpression is due to WAPL-dependent cohesin release, we deleted four amino acids (^1116^MEDC^1119^) that are critical for WAPL-dependent cohesin release (Fig. 8G) (*87, 88*). Although this “WAPL-dead” protein (WAPL^5AΔ4^) localized to nuclei, it had no effect on RAD21 foci following DNA damage (Fig. 8H, 8K). Interestingly, we also found that WAPL^5AΔ4^ overexpression caused nucleus-wide clustering of cohesin, similar to the “vermicelli” phenotype, although the cohesin threads appeared to be much thinner (Fig. 8H) (*21*), suggesting that this protein can act in a dominant negative fashion. Taken together, these results indicate that ATM promotes enrichment of cohesin by downregulating WAPL at damage foci in HeLa cells, and that the same mechanism promotes assembly of meiotic chromosome axes in response to nucleus-wide ATM activity.

## Discussion

We have found that downregulation of WAPL by ATM promotes cohesin localization along meiotic chromosome axes in *C. elegans* and at DNA repair foci in mammalian cells (Fig. 9). A key function of the axes is to regulate meiotic recombination (*89, 90*), which relies on DNA repair machinery, so it makes teleological sense that this assembly would be regulated by DDR signaling (*91*). Cohesins along the chromosome axis inhibit intersister repair and promote crossover formation between homologs (*20, 92–94*). The mechanisms that control axis assembly have been mysterious. While meiosis-specific cohesins are important, they are not sufficient, since meiotic cohesins can support mitosis to some degree (*95, 96*). Our findings illuminate the role of cohesin regulators, and how they are deployed during the unique cell cycle state of meiotic prophase.

**Fig. 9.**
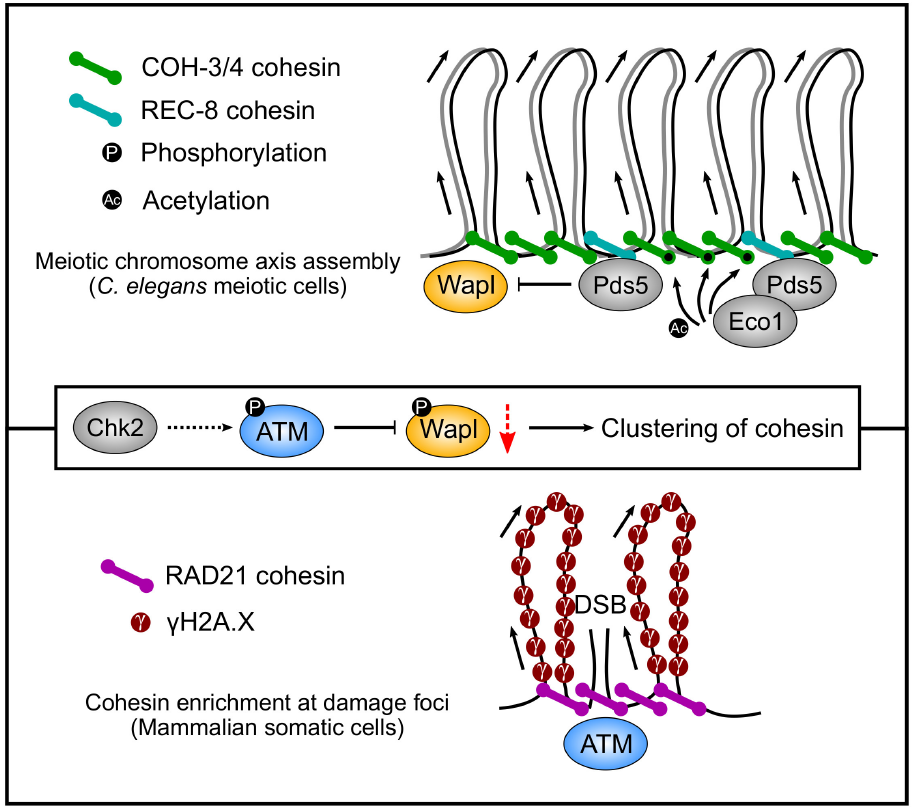
A conserved phosphoregulatory pathway in promoting cohesin clustering during meiosis and mitosis.

Nucleus-wide activation of ATM is important to regulate the formation of meiotic DSBs, and here we report that it also promotes axis assembly through down-regulation of WAPL, a key regulator of cohesin dynamics (*97, 98*). Previous studies have shown that WAPL-dependent cohesin release regulates cohesin-dependent loop extrusion (*21, 99*). Downregulation of WAPL is likely a conserved feature of meiosis, as loop anchors emerge along with axis compaction and the reduction of cohesin dynamics (*21, 99–101*). A role for ATM in meiotic axis morphogenesis was also demonstrated in *Arabidopsis* (*102*). Moreover, ATM and ATR localize sequentially to chromosome axes during early meiotic prophase in mice, consistent with roles in regulating cohesin (*103*). Our results also suggest that ATM downregulates WAPL at DNA damage foci in mitotic cells. Consistent with our observations, recent studies in humans show that ATM strengthens loop anchors following DNA damage (*104*), and persistent loop extrusion occurs at DNA damage foci (*105*).

Our findings also show that ECO-1 contributes to the stability of axial cohesins, although this was only apparent when downregulation of WAPL was defective. This is consistent with observations from *Drosophila* (*106*) and *Arabidopsis* (*54*). Our results suggest that ECO-1-dependent cohesin acetylation may also be regulated by CHK-2. In budding yeast, phosphorylation of the mitotic kleisin Mcd1 by Chk1 promotes Eco1-dependent Mcd1 acetylation, which in turn antagonizes Wapl and facilitates cohesion (*57, 107*). Analogous regulation of COH-3/4 by CHK-2 in *C. elegans* has also been proposed (*9*).

Spo76/Pds5 plays a widely conserved role in meiotic chromosome axis structure, and indeed is regarded as an axis component (*68*), but its specificity seems to vary across species. Studies from yeast to human have established the critical roles of Pds5 in regulating cohesin-dependent chromosomal events (*67, 69, 85, 108–112*). Our findings reveal that PDS-5 stabilizes REC-8 against WAPL, analogous to findings in fission yeast (*78*). However, in budding yeast and *Arabidopsis*, loss of Pds5 has little effect on the association of Rec8 with meiotic chromosomes (*74, 75, 113*). Nevertheless, budding yeast *PDS5* mutants show SC formation between sister chromatids rather than homologous chromosomes, a phenotype also seen in *C. elegans* and mouse spermatocytes lacking REC-8/Rec8 (*20, 74*).

The functional interplay between Pds5 and Wapl in different organisms has been enigmatic; in some cases these factors seem to act as a complex, while in others Pds5 antagonizes Wapl activity. Our results support a direct antagonism between *C. elegans* PDS-5 and WAPL-1 in meiotic axis assembly. We find that WAPL-1 can also function in the absence of PDS-5, and that PDS-5 and WAPL-1 localize to nuclei independently (data not shown). Most importantly, WAPL-1 depletion restores axial REC-8 cohesin even in the absence of PDS-5. Notably, the FGF and YSR motifs, which mediate binding of Wapl to Pds5 (*85, 86*), are both absent from *C. elegans* WAPL-1 (*64*). *C. elegans* PDS-5 also has a relatively long unstructured domain that may modulate its activities through mechanisms analogous to the function of Sororin in vertebrates.

Taken together, our work shows that axis assembly is driven by the specialized roles of meiotic cohesins and their interactions with cohesin regulators, which in turn are controlled by the unique DDR signaling during meiotic prophase.

## Acknowledgments

We thank Douglas Koshland, Vincent Guacci, Siheng Xiang, Lorenzo Costantino and Kevin Boardman, and all Dernburg lab members for insightful discussions throughout this study and for critical reading of the manuscript. We also thank Weiguo Zhang for help with the mammalian cell experiments, and Christina Glazier for preliminary analysis of WAPL-1 function in *C. elegans*.

## Author contributions

Conceptualization: ZY, AFD; Methodology: ZY, AFD; Investigation: ZY, AFD; Visualization: ZY, AFD; Funding acquisition: AFD; Project administration: AFD; Supervision: AFD; Writing – original draft: ZY; Writing – review & editing: ZY, AFD.

## Data and materials availability

All data are available in the main text or the supplementary materials. Additional materials/information will be provided upon request.

## Materials and Methods

### Strain maintenance

All *C. elegans* strains were maintained on standard nematode growth medium (NGM) plates seeded with OP50 bacteria at 20°C (*114*). Young adults (20-24 hrs post-L4) were used for immunofluorescence analysis.

### Strain construction

All new alleles used in this study were generated by CRISPR/Cas9-mediated genome editing. See Table S1 for the list of strains used in this study. Briefly, Alt-R®CRISPR-Cas9 crRNAs specific for target sites were mixed with a *dpy-10*-specific crRNA (*115*)) at a molar ratio of 8:1. These were denatured and annealed to an equal quantity of tracrRNA (Integrated DNA Technologies, Coralville, IA) by heating to 95°C for 5 minutes, followed by 5 minutes at 25°C. 1 μL of 100μM hybridized tracrRNA/crRNA was combined with 2.5 μL of 40 μM *S. pyogenes* Cas9-NLS purified protein (QB3 MacroLab, UC Berkeley, Berkeley, CA) and incubated at room temperature for 5 minutes. 0.5 μL of 100 μM stock of an Ultramer® DNA oligo (IDT) repair template containing 35-45 bp homology arms and the desired epitope/degron or mutation sequence was added to the mixture, for a total volume of 5 μL, and injected into the gonads of young adult hermaphrodites aged 24 hours from the late L4 stage. Injected P_0_ animals were maintained on individual plates at 20°C for 3-4 days. Roller and Dumpy F1 were singled, maintained at 20°C for 3 days, and screened by PCR for the desired mutation or epitope tag. Candidate alleles were verified by Sanger sequencing. See Table S2 for a complete list of crRNA, repair template, and genotyping primer sequences used in this study.

### Worm viability and fertility

To quantify brood sizes, male self-progeny, and embryonic viability, L4 hermaphrodites were plated individually and transferred to new plates daily for four consecutive days. Eggs were counted twice a day to minimize counting errors. Viable progeny and males were scored when they reached young adulthood.

### Auxin-induced protein depletion in worms

Auxin-induced depletion of degron-tagged proteins was performed as previously described (*116*). Unless otherwise indicated,hermaphrodites at the L4 stage were transferred to seeded plates containing 1 mM indole-3-acetic acid (IAA, auxin) and incubated for 24 hrs before analysis.

### Plasmids

To express human WAPL in HeLa cells, sequences were inserted into the pcDNA3-acGFP vector, obtained from Addgene (Cat No. 128047). The WAPL coding sequence was divided into four ~1 kb fragments and synthesized by Twist Bioscience. These fragments were inserted at the 3’ end of the GFP coding sequence using Gibson assembly (*117*) and verified by Sanger sequencing.

### Antibodies and reagents

Primary antibodies were purchased from commercial sources or have been described in previous studies, and were diluted as follows: rabbit anti-RAD-51 (1:500, (*30*)), rabbit anti-pHIM-8/ZIMs (1:500, (*27*)), goat anti-SYP-1 (1:300, (*30*)), chicken anti-HTP-3 (1:500, (*118*)), mouse anti-HA (1:400, Thermo Fisher 26183), mouse anti-FLAG (1:500, Sigma F3165), mouse anti-V5 (1:500, Thermo Fisher R960-25), rabbit anti-V5 (1:250, Millipore Sigma V8137), mouse anti-WAPL (1:500, Santa Cruz sc-365189), rabbit anti-γH2A.X antibody (1:500, Cell Signaling, Cat No. 2577), mouse anti-ATM antibody (1:500, Thermo Fisher, Cat No. MA1-23152), rabbit anti-pS/TQ antibody (1:500,Cell Signaling,Cat No. 6966), rabbit anti-COH-3/4 antibody (1:500,SDQ3972, ModENCODE project (*119*)), rabbit anti-REC-8 antibody (1:500, SDQ0802, ModENCODE project (*119*)), rabbit anti-WAPL-1 antibody (1:500, SDQ3963, ModENCODE project (*119*)). Secondary antibodies raised in donkey and labeled with Alexa 488, Cy3, or Cy5 (Jackson ImmunoResearch Laboratories) and used at 1:400 dilution. Kinase inhibitors included VE-821 (Selleckchem S8007); NU7441 (Selleckchem S2638); and KU-55933 (Selleckchem S1092). DNA damage-inducing agents included etoposide (Sigma Cat No. E1383) and bleomycin (Fisher Cat No. B397210MG).

### siRNA-mediated knockdown

The following ON-TARGETplus SMARTpool siRNAs were purchased from Horizon Discovery: control pool, Cat No. D-001810-10-05, WAPL siRNA, Cat No. L-026287-01-0005, ATM siRNA, Cat No. Cat No. L-003201-00-0005. ATR siRNA, Cat No. L-003202-00-0005. HeLa cells were cultured on coverslips in 6-well plates to 25% confluency, and siRNA knock-down was performed using DharmaFECT™ according to the manufacturer’s recommendations. Cells were fixed and analyzed 72 hrs after siRNA transfection. DNA damage-inducing agents and/or kinase inhibitors were added 24 hrs before fixation.

### Transient transfection

For WAPL overexpression, HeLa cells were grown on coverslips in 6-well plates to 50% confluency. 2.5 μg of purified plasmid DNA was mixed with 5 μl Lipofectamine 3000 (Thermo Fisher) and used for transfection according to the manufacturer’s protocol. Cells were fixed for imaging 48 hrs post transfection. DNA damage-inducing agents and/or kinase inhibitors were added 24 hrs before fixation.

### Chemical treatments

DNA damage was induced by addition of 0.8 μM etoposide or 0.4 μM bleomycin for 24 hrs. KU-55933, VE-821 and NU7441 added at 1 μM for 24 hrs.

### Immunofluorescence assays

Adult hermaphrodites were dissected on a clean coverslip in egg buffer (25 mM HEPES pH 7.4, 118 mM NaCl, 48 mM KCl, 2 mM EDTA, 0.5 mM EGTA) containing 0.01% tetramisole and 0.1% Tween-20. Samples were fixed for 2 minutes in egg buffer containing 1% formaldehyde and then transferred to a 1.5 mL tube containing PBS + 0.1% Tween-20 (PBST). After 5 minutes, the buffer was replaced with ice-cold methanol and incubated at −20°C for an additional 10 minutes. Worms were washed twice with PBST, blocked with Roche blocking reagent diluted into PBST, and stained with primary antibodies diluted in blocking solution at 4°C overnight. Samples were then washed with PBST and incubated with secondary antibodies diluted in blocking solution at room temperature for 1 hr. Worms were washed twice with PBST and mounted in ProLong™ Diamond with DAPI (Invitrogen) before imaging.

For immunofluorescence of HeLa cells, coverslips in 6-well plates were washed with PBS and then fixed with 4% formaldehyde in PBS at room temperature for 10 min. After 3 washes with PBS, cells were permeabilized by addition of 0.5% Triton X-100 in PBS at room temperature for 5 min. They were rinsed with PBS and blocked with 5% BSA in PBS at room temperature for 1 hr. Cells were then washed with PBS and incubated with primary antibodies diluted in 1% BSA at room temperature for 2 hrs. After another PBS wash, cells were incubated in secondary antibodies diluted in 1% BSA in PBS at room temperature for 1 hr in dark. Cells were then washed again with PBS and mounted in ProLong™ Diamond with DAPI before imaging.

### Microscopy

All images were acquired as z-stacks of optical sections at 0.2 μm intervals using a DeltaVision Elite microscope (GE) with a 100x 1.4 N.A. or 60x 1.42 N.A. oil-immersion objective. Iterative 3D deconvolution, image projection, and colorization were performed using the softWoRx package, ImageJ and Adobe Photoshop CC 2017, respectively.

### Image analysis

To quantify the abundance of proteins in *C. elegans* germline nuclei, additive projections were generated from raw (undeconvolved) 3D data stacks after background subtraction using the rolling ball tool in ImageJ. Individual nuclei (regions of interest, ROI) were manually segmented based on DAPI staining in ImageJ, and the integrated intensity within each ROI was calculated. For each condition, 80-200 nuclei from 2-4 gonads were quantified.

To quantify protein abundance in HeLa cell nuclei, individual nuclei (ROIs) were first segmented based on DAPI fluorescence in an equatorial optical section from a 3D image stack using the 2D watershed tool. Protein abundance (integrated intensity) within this region was calculated from additive Z projections, similar to the approach we used to quantify proteins in *C. elegans* germline nuclei.

To quantify RAD21 enrichment at sites of DNA damage in HeLa cells, we developed an automated method to ensure consistency and minimize potential investigator bias. Following empirical optimization, the method was applied to each dataset using ImageJ macros. For experiments involving expression of GFP or GFP-WAPL, only GFP-positive cells were included; these were identified based on GFP fluorescence in equatorial sections using the Auto Threshold tool in ImageJ in “Li” maximum entropy mode. Nuclear ROIs were segmented as described above. Peaks of immunofluorescence of RAD21 and DNA damage markers (γH2A.X or pS/TQ) were segmented using the Auto Threshold tool in MaxEntropy mode. Each optical section within 3D image stacks spanning the nuclear volume was analyzed. RAD21 regions smaller than 0.1 μm^2^ or detected in only a single optical section were discarded as unreliable. RAD21-positive peaks were also rejected if <50% of their area overlapped with DNA damage markers. The resulting 3D binary masks of RAD21-enriched regions were projected to create a single image, then labeled and counted using the measure module of the scikit-image library in Python.

### Data presentation

For data based on immunofluorescence in *C. elegans* germline nuclei, we show representative images of the distal regions of dissected gonads. All images are oriented with the distal tip on the left. They show the entire proliferative (premeiotic) region and a similarly-sized region containing nuclei in early meiotic prophase. The boundary between premeiotic and meiotic prophase is indicated by a dashed line. Figure labels indicate proteins that were depleted by auxin treatment. For immunofluorescence in HeLa cells, representative images of individual nuclei are shown with enlargements of fluorescent foci as insets.

For quantitative analysis of immunofluorescence, the integrated nuclear intensity or number of foci were measured as described above under “Image analysis.” Tukey boxplots of data points from individual nuclei were generated using R. Boxes indicate the quartiles and median, and the median value is also indicated next to the box. The number of nuclei that were scored for each condition/group is shown in parentheses underneath the data points.

### Statistical analysis

Unless otherwise specified, data from different conditions were compared using Student’s *t*-test. The number of asterisks indicates the calculated *p*-values: n.s. (not significant) p≥0.05, \**p*<0.05, \*\**p*<0.01, \*\*\**p*<0.001, \*\*\*\**p*<0.0001.

## Supplementary Text

### The activity of *C. elegans* ATL-1 is independent of phosphorylation at a CHK-2 motif

Like *C. elegans* ATM-1, ATL-1 also has a CHK-2 consensus at the C-terminus of its FAT domain (Fig. S3A). To test whether this CHK-2 consensus motif plays a similar role in regulating ATL-1, we constructed a non-phosphorylatable mutant (*atl-1^KA^*) and assayed the embryonic viability and meiotic nondisjunction of homozygous mutant animals (*120*). As previously reported, the *atl-1* null allele *tm853* showed 100% embryonic lethality (Fig. S3B) (*121*). By contrast, animals homozygous for the *atl-1^KA^* allele showed normal viability and fertility, indicating that this motif is dispensable for ATL-1 activity (Fig. S3B).

### WAPL depletion leads to the formation of “vermicelli” in the absence of ATM

Our results indicate that ATM downregulates WAPL to promote cohesin enrichment at DNA damage foci. We therefore wondered whether depletion of WAPL might restore RAD21 foci upon DNA damage in the absence of ATM activity. However, WAPL knockdown resulted in the appearance of “vermicelli” even in the absence of ATM (Fig. S7I-S7K), likely reflecting global stabilization of cohesin. Under these conditions, we did not detect any specific enrichment of cohesin at damage foci.

**Fig. S1.**
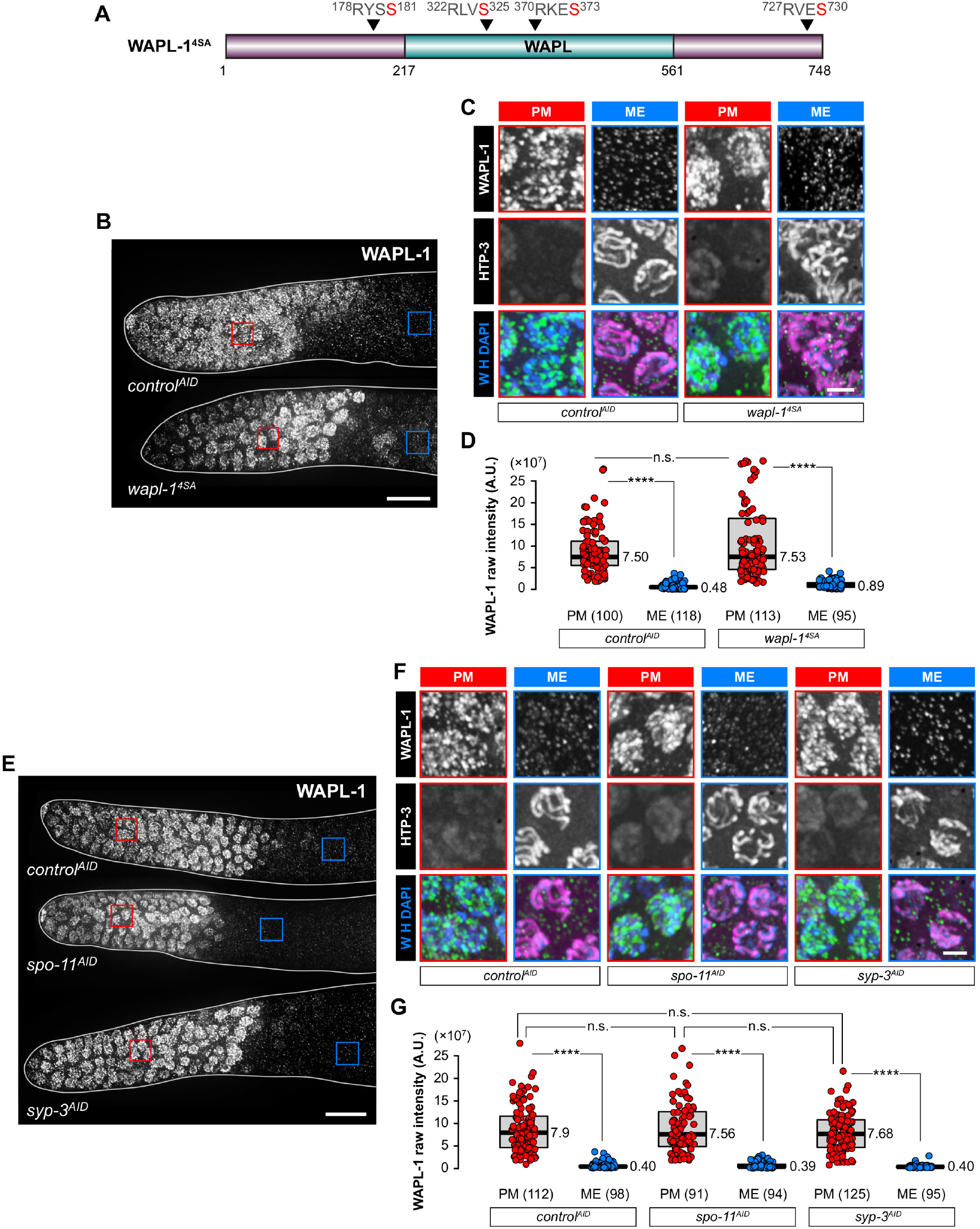
WAPL-1 downregulation does not require direct phosphorylation by CHK-2, DSBs, or synapsis. **(A)** WAPL-1 sequence indicating the positions of CHK-2 consensus motifs that were mutated in the *wapl-1^4SA^* allele. **(B)** and **(E)** WAPL-1 immunostaining germline nuclei. Scale bar, 10 μM. **(C)** and **(F)** Enlarged images of the regions outlined in (B) and (E). HTP-3 marks chromosome axes in meiotic prophase nuclei. Scale bar, 2 μM. **(D)** and **(G)** Quantification of WAPL-1 intensity in (B) and (E), respectively.

**Fig. S2.**
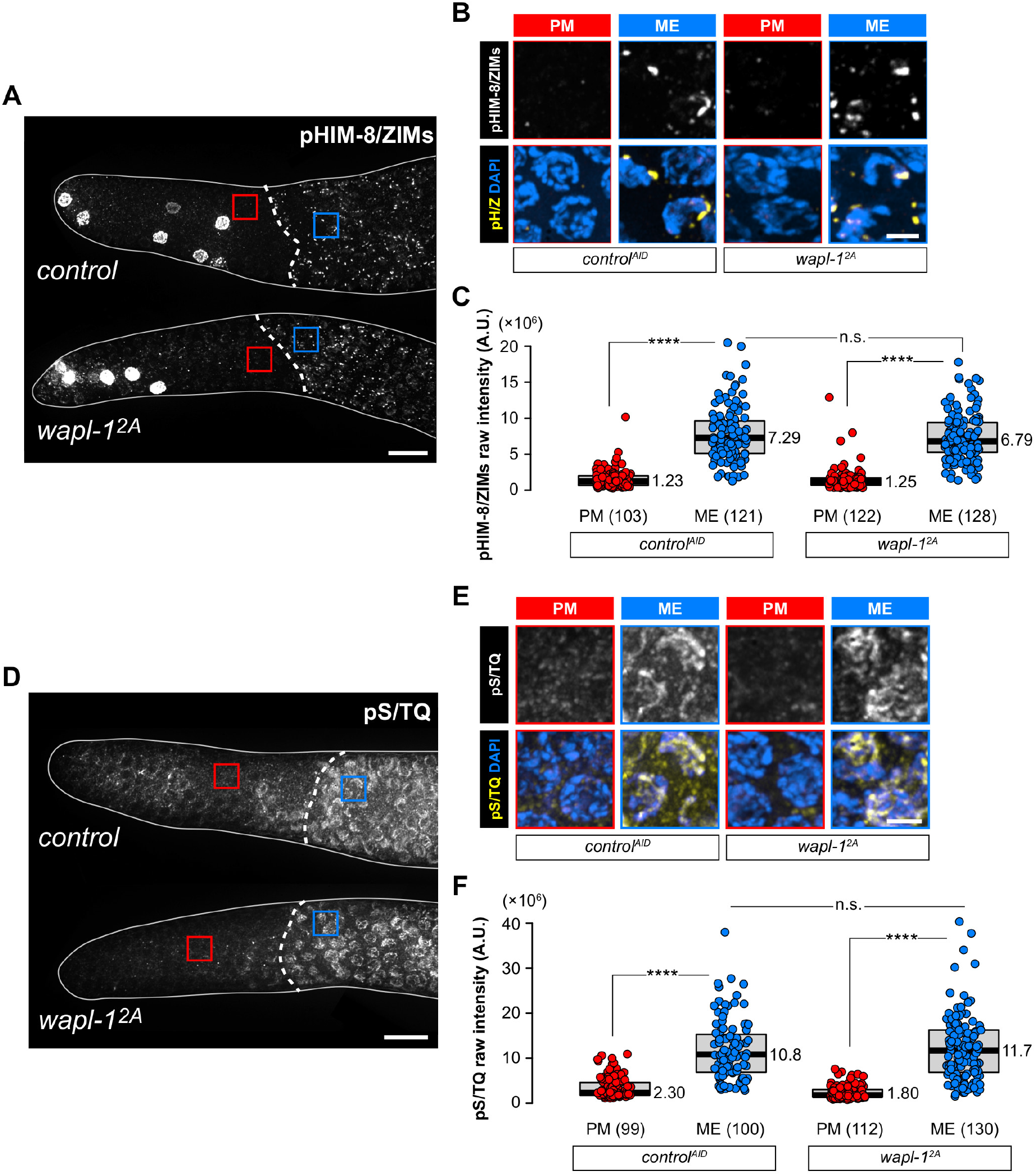
Mutation of ATM target sites in WAPL-1 does not affect CHK-2 or ATM-1 activity at meiotic entry. **(A)** and **(D)** pHIM-8/ZIM (A) and pS/TQ (D) immunostaining in the distal tip of gonads of wild-type and *wapl-1^2A^* worms. The pHIM-8/ZIM antibody also recognizes an unidentified, CHK-2-independent epitope in mitotic cells (*27*). Scale bar, 10 μM. **(B)** and **(E)** Enlarged images of the regions outlined in (A) and (D), respectively. Scale bar, 2 μM. **(C)** and **(F)** Quantification of the intensity of pHIM-8/ZIMs (C) and pS/TQ (F).

**Figure S3.**
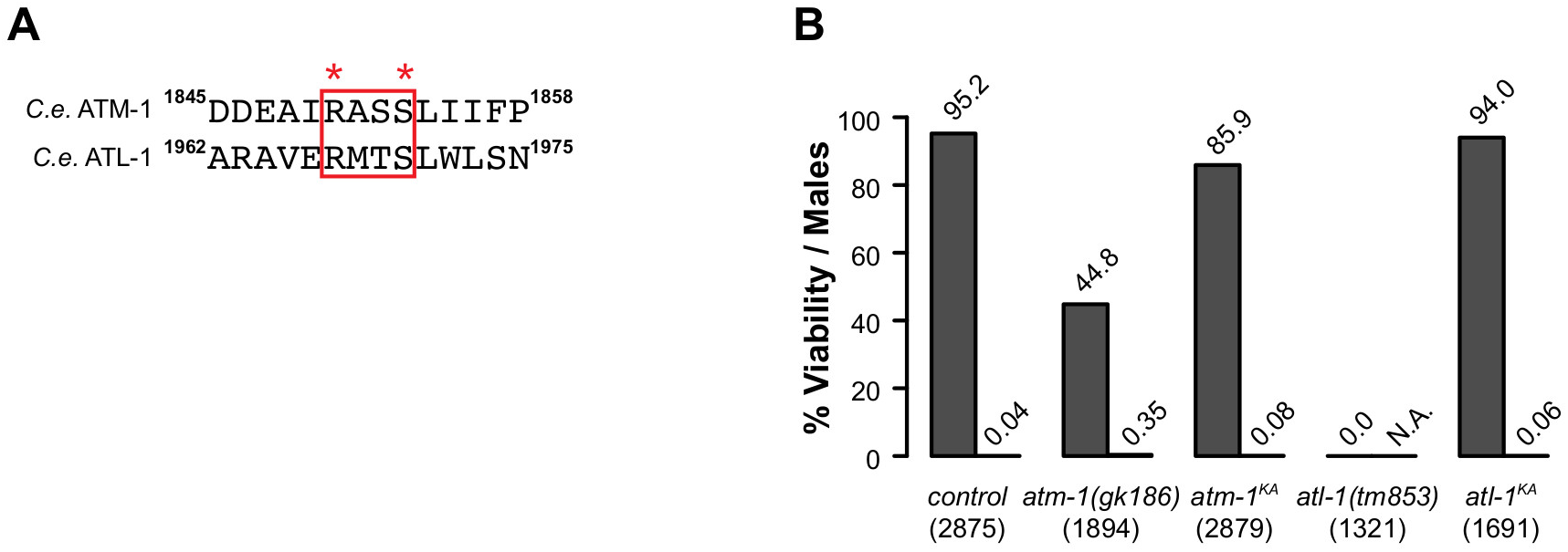
A putative CHK-2 target site in the FAT domain of ATL-1 does not promote its kinase activity. **(A)** Alignment of residues at the C-terminus of the FAT domains of *C. elegans* ATM-1 and ATL-1. CHK-2 consensus motifs are highlighted. The conserved phospho-serines and arginines at the −3 position are marked by asterisks. **(B)** Viability of embryos and frequency of male self-progeny from hermaphrodites homozygous for the indicated alleles. Null alleles of both *atm-1* (*atm-1(gk186)*) and *atl-1* (*atl-1(tm853)*) were assayed for comparison. The number of eggs scored for each allele is shown in parentheses.

**Figure S4.**
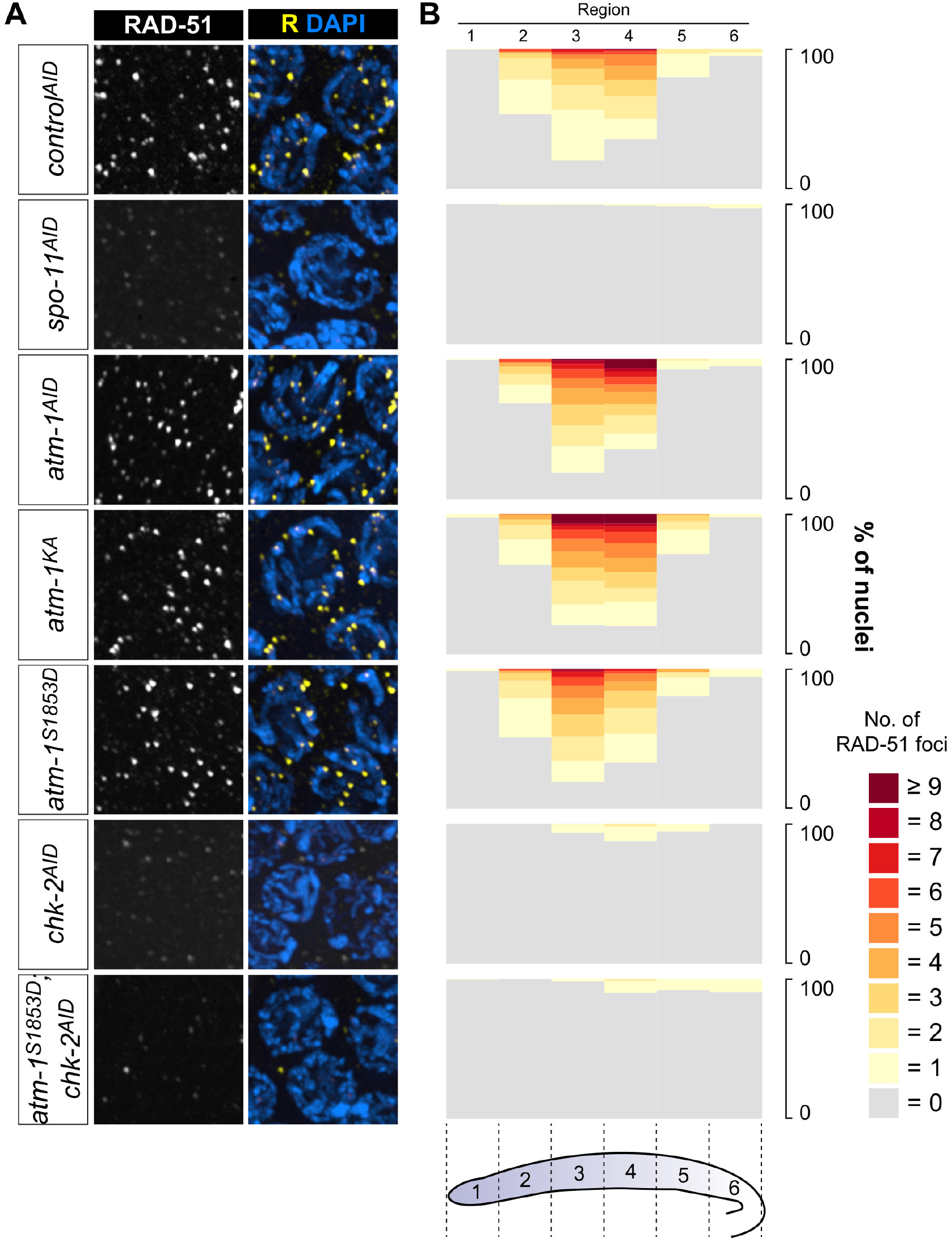
DSBs are dispensable for a basal level of ATM-1 activity in meiotic prophase nuclei. **(A)** RAD-51 foci indicate the presence or absence of meiotic DSBs in pachytene nuclei. Scale bar, 2 μM. **(B)** Quantitative analysis of RAD-51 foci. The region of the germline from the distal tip to late pachytene was divided into six zones of equal length. The distribution of RAD-51 foci per nucleus nuclei for each region is shown for each of the conditions represented by images in (A).

**Figure S5.**
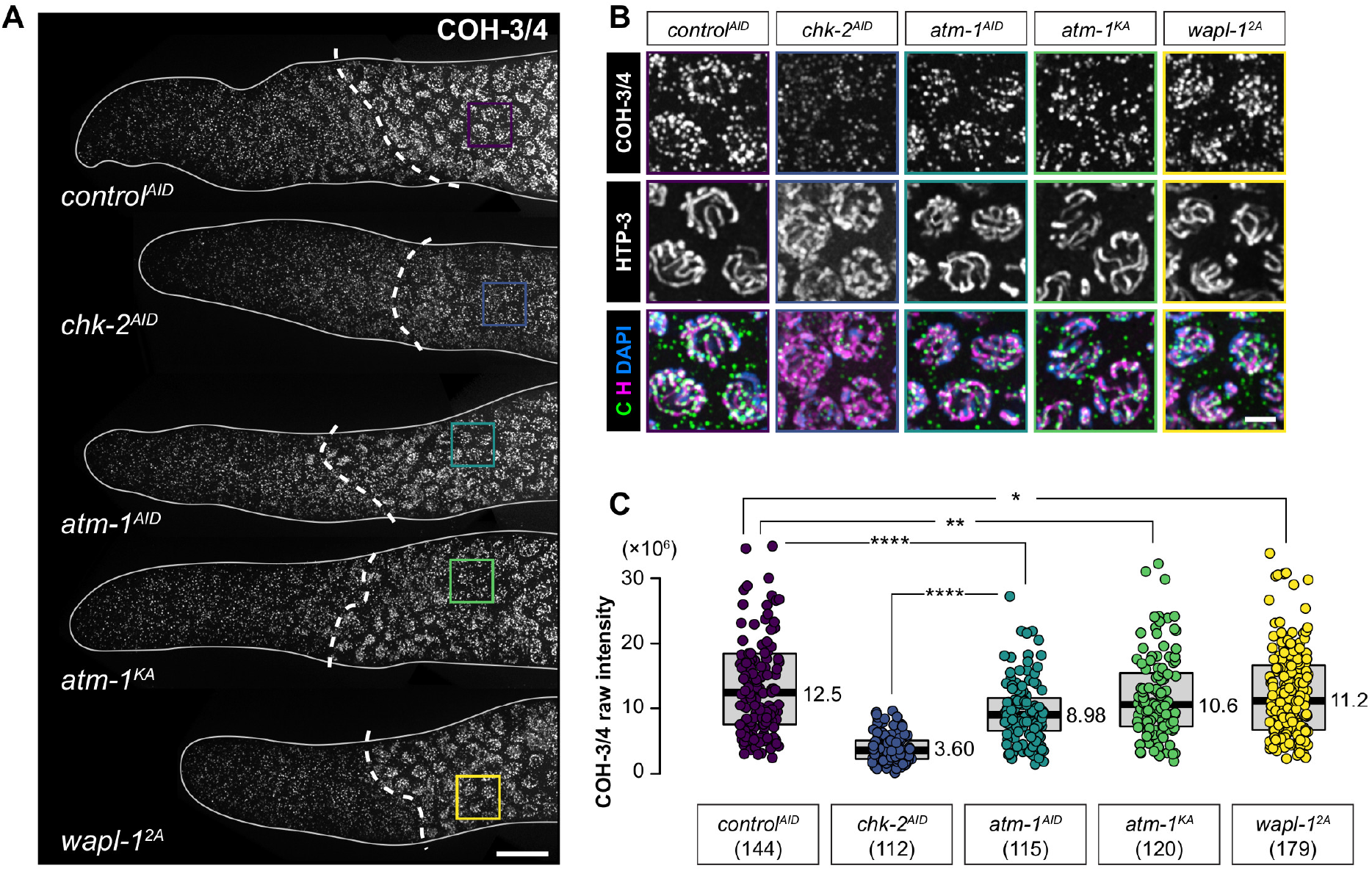
CHK-2 promotes axial cohesin stabilization. **(A)** COH-3/4 immunofluorescence in the distal region of gonads. Dashed lines indicate the boundaries between premeiotic and meiotic germline. Scale bar, 10 μM. **(B)** Enlarged images of the regions indicated in (A). HTP-3 immunostaining (in magenta) marks chromosome axes. Scale bar, 2 μM. **(C)** Quantification of the intensity of COH-3/4 immunostaining in (A).

**Figure S6.**
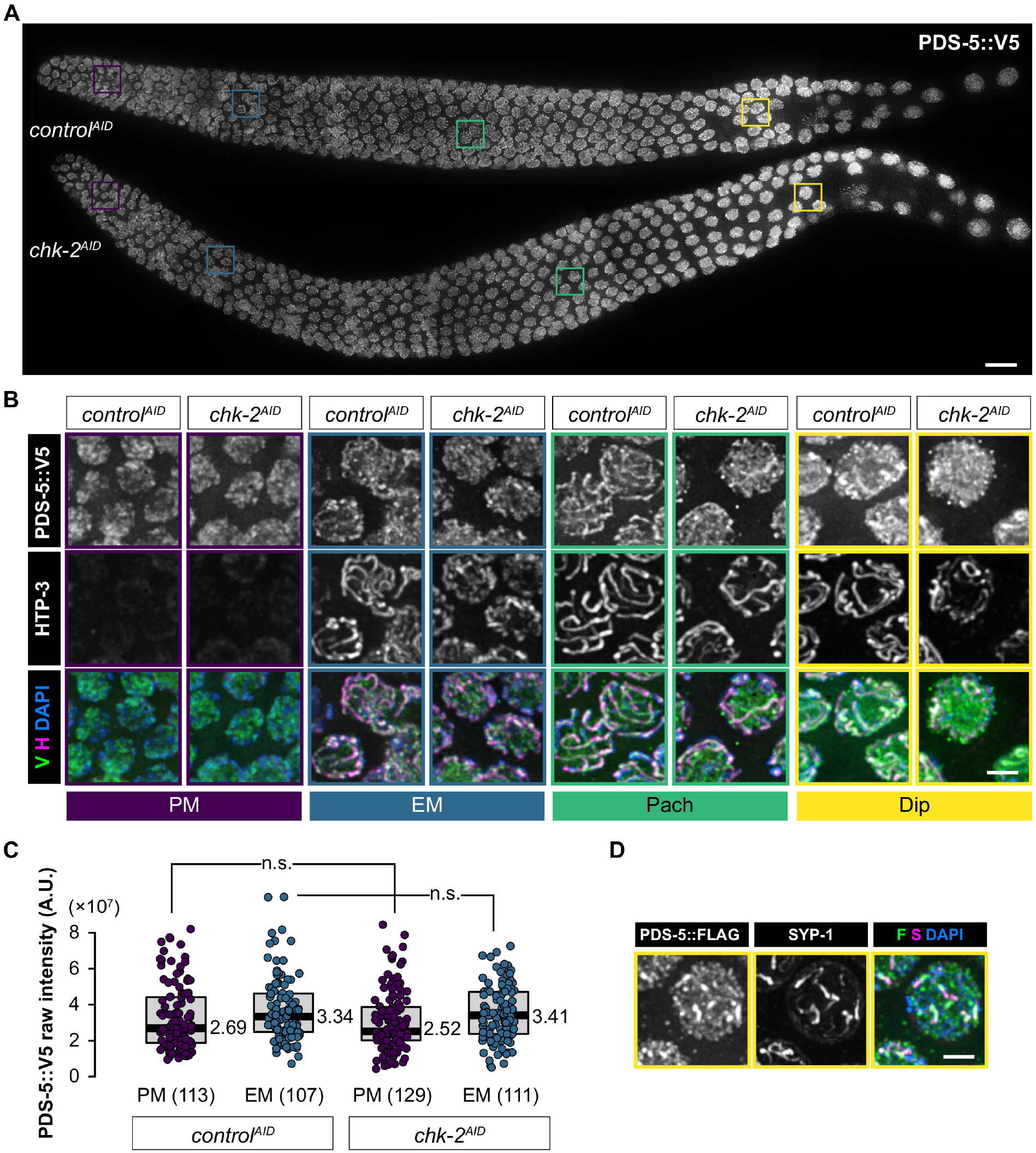
PDS-5 localizes to meiotic chromosome axes independently of CHK-2. **(A)** PDS-5::V5 immunofluorescence using anti-V5 in *C. elegans* gonads. Scale bar, 10 μM. **(B)** Enlarged images of the regions outlined in (A). HTP-3 immunostaining (in magenta) marks chromosome axes. Scale bar, 2 μM. **(C)** Quantification of the PDS-5::V5 intensity in (B). **(D)** PDS::FLAG immunofluorescence in diplotene nuclei. SYP-1 immunofluorescence in late diplotene nuclei is restricted to the “short arm” of each bivalent. Scale bar, 2 μM.

**Figure S7.**
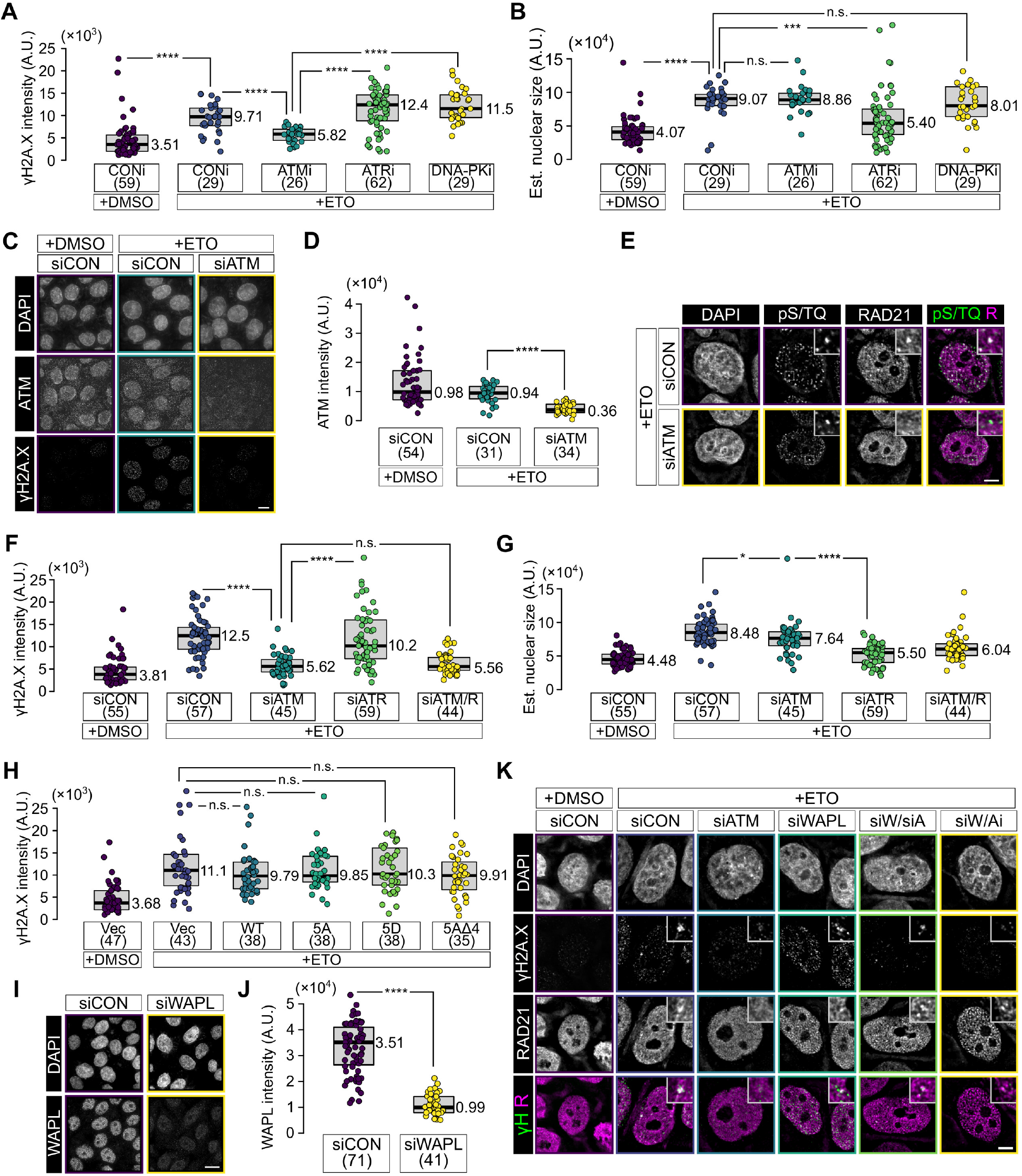
ATM-mediated WAPL downregulation promotes cohesin enrichment at DNA damage foci. **(A)** and **(B)** Quantification of nuclear γH2A.X intensity (A) and nuclear size (B) in cells that were exposed to etoposide (ETO)-induced following chemical inhibition of DNA damage kinases. ATM was inhibited by addition of KU55933 (ATMi), ATR by VE-821 (ATRi), and DNA-PK by NU7441 (DNA-Pki). **(C)** Immunofluorescence of ATM in HeLa cells treated with control siRNA or siRNA targeting ATM, with or without etoposide (ETO) treatment. Both ATM immunostaining and γH2A.X immunostaining were used to assess ATM knock-down efficiency. Scale bar, 10 μM. **(D)** Quantification of nuclear ATM intensity in cells treated as in (C). Integrated ATM intensity was normalized against DAPI intensity for each nucleus. **(E)** Immunofluorescence of RAD21 in nuclei of HeLa cells treated with either control siRNA or siRNA against ATM, and then etoposide (ETO) to induce DNA damage. DNA damage foci are marked by pS/TQ immunofluorescence. Scale bar, 10 μM. **(F)** and **(G)** Quantification of nuclear γH2A.X intensity (F) and estimated nuclear size (G) in cells following siRNA-mediated knockdown of ATM and/or ATR. The integrated γH2A.X intensities were normalized by integrated DAPI intensities for each nucleus. **(H)** Quantification of nuclear γH2A.X intensity in transfected HeLa cells expressing GFP or GFP-WAPL fusion proteins, following etoposide (ETO)-induced DNA damage. **(I)** Immunofluorescence of WAPL in nuclei of HeLa cells treated with either control siRNA or siRNA against WAPL. Scale bar, 10 μM. **(J)** Quantification of nuclear WAPL intensity in HeLa cells, normalized against DAPI intensity in each nucleus (I). **(K)** Immunofluorescence of RAD21 in nuclei of HeLa cells treated with the indicated siRNA and then etoposide (ETO). WAPL siRNA was combined with KU55933 (ATMi) or siRNA targeting ATM. γH2A.X immunofluorescence marks DNA damage foci. Scale bar, 10 μM.

**Table S1.**
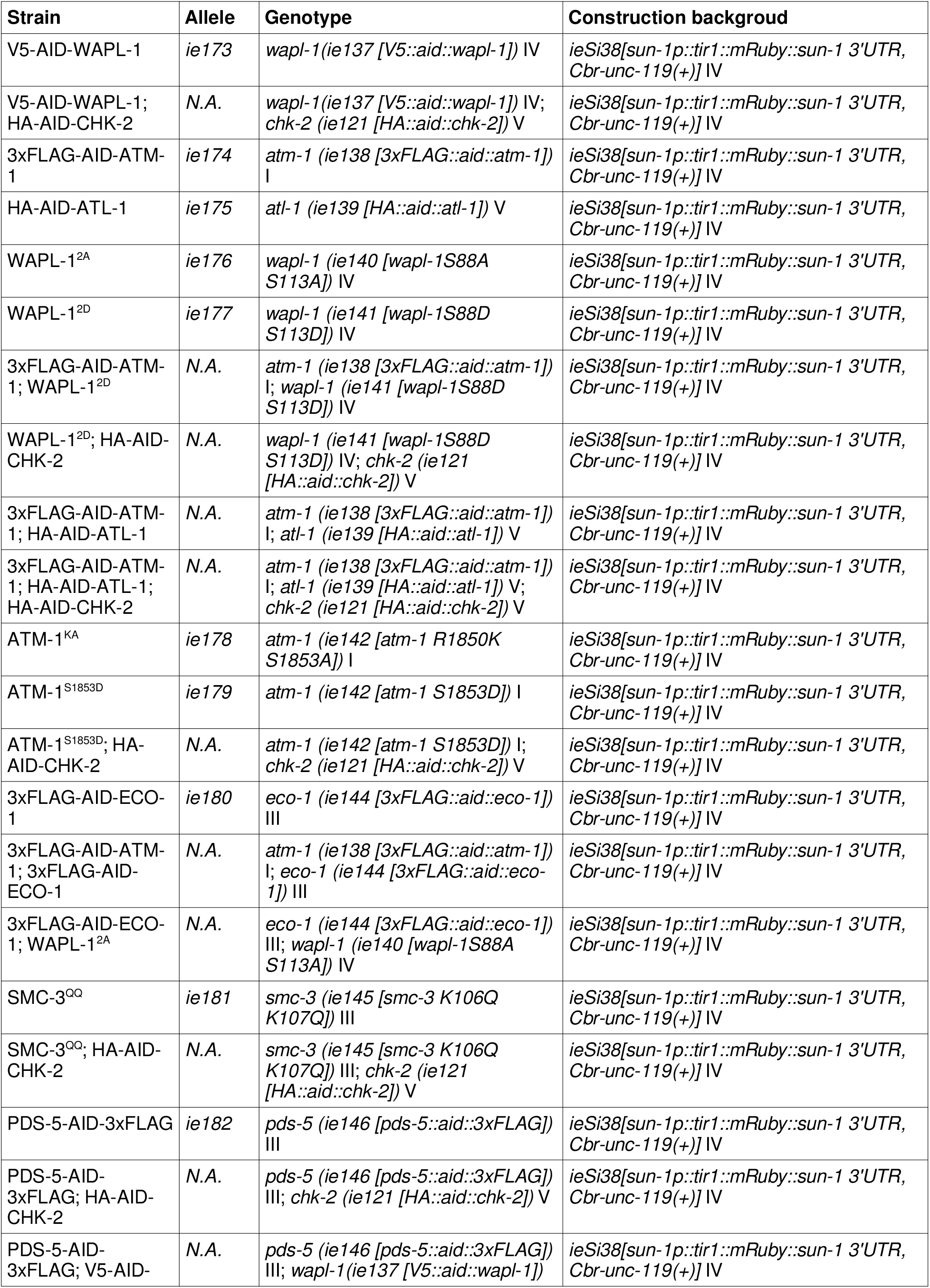
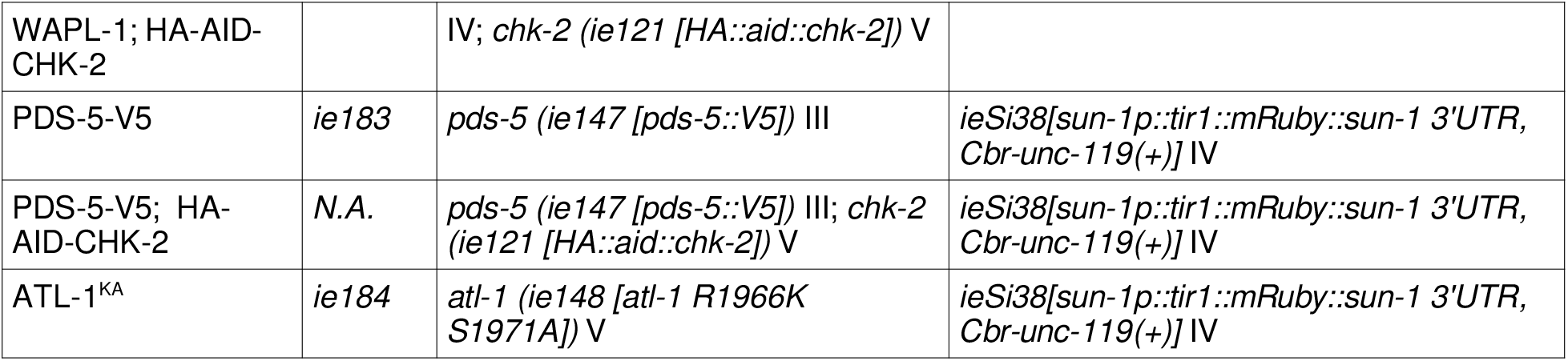
Strains generated in this study.

| **Strain** | **Allele** | **Genotype** | **Construction background** |
| --- | --- | --- | --- |
| V5-AID-WAPL-1 | *ie173* | *wapl-1* (*ie137* [*V5::aid::wapl-1*]) IV | *ieSi38*[*sun-1p::tir1::mRuby::sun-1 3'UTR*, *Cbr-unc-119(+)*] IV |
| V5-AID-WAPL-1;<br>HA-AID-CHK-2 | N.A. | *wapl-1* (*ie137* [*V5::aid::wapl-1*]) IV;<br>*chk-2* (*ie121* [*HA::aid::chk-2*]) V | *ieSi38*[*sun-1p::tir1::mRuby::sun-1 3'UTR*, *Cbr-unc-119(+)*] IV |
| 3xFLAG-AID-ATM-1 | *ie174* | *atm-1* (*ie138* [*3xFLAG::aid::atm-1*]) I | *ieSi38*[*sun-1p::tir1::mRuby::sun-1 3'UTR*, *Cbr-unc-119(+)*] IV |
| HA-AID-ATL-1 | *ie175* | *atl-1* (*ie139* [*HA::aid::atl-1*]) V | *ieSi38*[*sun-1p::tir1::mRuby::sun-1 3'UTR*, *Cbr-unc-119(+)*] IV |
| WAPL-1<sup>2A</sup> | *ie176* | *wapl-1* (*ie140* [*wapl-1S88A* *S113A*]) IV | *ieSi38*[*sun-1p::tir1::mRuby::sun-1 3'UTR*, *Cbr-unc-119(+)*] IV |
| WAPL-1<sup>2D</sup> | *ie177* | *wapl-1* (*ie141* [*wapl-1S88D* *S113D*]) IV | *ieSi38*[*sun-1p::tir1::mRuby::sun-1 3'UTR*, *Cbr-unc-119(+)*] IV |
| 3xFLAG-AID-ATM-1;<br>WAPL-1<sup>2D</sup> | N.A. | *atm-1* (*ie138* [*3xFLAG::aid::atm-1*]) I;<br>*wapl-1* (*ie141* [*wapl-1S88D* *S113D*]) IV | *ieSi38*[*sun-1p::tir1::mRuby::sun-1 3'UTR*, *Cbr-unc-119(+)*] IV |
| WAPL-1<sup>2D</sup>; HA-AID-CHK-2 | N.A. | *wapl-1* (*ie141* [*wapl-1S88D* *S113D*]) IV; *chk-2* (*ie121* [*HA::aid::chk-2*]) V | *ieSi38*[*sun-1p::tir1::mRuby::sun-1 3'UTR*, *Cbr-unc-119(+)*] IV |
| 3xFLAG-AID-ATM-1;<br>HA-AID-ATL-1 | N.A. | *atm-1* (*ie138* [*3xFLAG::aid::atm-1*]) I;<br>*atl-1* (*ie139* [*HA::aid::atl-1*]) V | *ieSi38*[*sun-1p::tir1::mRuby::sun-1 3'UTR*, *Cbr-unc-119(+)*] IV |
| 3xFLAG-AID-ATM-1;<br>HA-AID-ATL-1;<br>HA-AID-CHK-2 | N.A. | *atm-1* (*ie138* [*3xFLAG::aid::atm-1*]) I;<br>*atl-1* (*ie139* [*HA::aid::atl-1*]) V;<br>*chk-2* (*ie121* [*HA::aid::chk-2*]) V | *ieSi38*[*sun-1p::tir1::mRuby::sun-1 3'UTR*, *Cbr-unc-119(+)*] IV |
| ATM-1<sup>KA</sup> | *ie178* | *atm-1* (*ie142* [*atm-1 R1850K* *S1853A*]) I | *ieSi38*[*sun-1p::tir1::mRuby::sun-1 3'UTR*, *Cbr-unc-119(+)*] IV |
| ATM-1<sup>S1853D</sup> | *ie179* | *atm-1* (*ie142* [*atm-1 S1853D*]) I | *ieSi38*[*sun-1p::tir1::mRuby::sun-1 3'UTR*, *Cbr-unc-119(+)*] IV |
| ATM-1<sup>S1853D</sup>; HA-AID-CHK-2 | N.A. | *atm-1* (*ie142* [*atm-1 S1853D*]) I;<br>*chk-2* (*ie121* [*HA::aid::chk-2*]) V | *ieSi38*[*sun-1p::tir1::mRuby::sun-1 3'UTR*, *Cbr-unc-119(+)*] IV |
| 3xFLAG-AID-ECO-1 | *ie180* | *eco-1* (*ie144* [*3xFLAG::aid::eco-1*]) III | *ieSi38*[*sun-1p::tir1::mRuby::sun-1 3'UTR*, *Cbr-unc-119(+)*] IV |
| 3xFLAG-AID-ATM-1;<br>3xFLAG-AID-ECO-1 | N.A. | *atm-1* (*ie138* [*3xFLAG::aid::atm-1*]) I;<br>*eco-1* (*ie144* [*3xFLAG::aid::eco-1*]) III | *ieSi38*[*sun-1p::tir1::mRuby::sun-1 3'UTR*, *Cbr-unc-119(+)*] IV |
| 3xFLAG-AID-ECO-1;<br>WAPL-1<sup>2A</sup> | N.A. | *eco-1* (*ie144* [*3xFLAG::aid::eco-1*]) III;<br>*wapl-1* (*ie140* [*wapl-1S88A* *S113A*]) IV | *ieSi38*[*sun-1p::tir1::mRuby::sun-1 3'UTR*, *Cbr-unc-119(+)*] IV |
| SMC-3<sup>QQ</sup> | *ie181* | *smc-3* (*ie145* [*smc-3 K106Q* *K107Q*]) III | *ieSi38*[*sun-1p::tir1::mRuby::sun-1 3'UTR*, *Cbr-unc-119(+)*] IV |
| SMC-3<sup>QQ</sup>; HA-AID-CHK-2 | N.A. | *smc-3* (*ie145* [*smc-3 K106Q* *K107Q*]) III; *chk-2* (*ie121* [*HA::aid::chk-2*]) V | *ieSi38*[*sun-1p::tir1::mRuby::sun-1 3'UTR*, *Cbr-unc-119(+)*] IV |
| PDS-5-AID-3xFLAG | *ie182* | *pds-5* (*ie146* [*pds-5::aid::3xFLAG*]) III | *ieSi38*[*sun-1p::tir1::mRuby::sun-1 3'UTR*, *Cbr-unc-119(+)*] IV |
| PDS-5-AID-3xFLAG;<br>HA-AID-CHK-2 | N.A. | *pds-5* (*ie146* [*pds-5::aid::3xFLAG*]) III;<br>*chk-2* (*ie121* [*HA::aid::chk-2*]) V | *ieSi38*[*sun-1p::tir1::mRuby::sun-1 3'UTR*, *Cbr-unc-119(+)*] IV |
| PDS-5-AID-3xFLAG;<br>V5-AID- | N.A. | *pds-5* (*ie146* [*pds-5::aid::3xFLAG*]) III;<br>*wapl-1* (*ie137* [*V5::aid::wapl-1*]) IV | *ieSi38*[*sun-1p::tir1::mRuby::sun-1 3'UTR*, *Cbr-unc-119(+)*] IV |
| WAPL-1; HA-AID-CHK-2 |  | IV; *chk-2* (*ie121* [*HA::aid::chk-2*]) V |  |
| PDS-5-V5 | *ie183* | *pds-5* (*ie147* [*pds-5::V5*]) III | *ieSi38*[*sun-1p::tir1::mRuby::sun-1 3'UTR*, *Cbr-unc-119(+)*] IV |
| PDS-5-V5; HA-AID-CHK-2 | N.A. | *pds-5* (*ie147* [*pds-5::V5*]) III; *chk-2* (*ie121* [*HA::aid::chk-2*]) V | *ieSi38*[*sun-1p::tir1::mRuby::sun-1 3'UTR*, *Cbr-unc-119(+)*] IV |
| ATL-1<sup>KA</sup> | *ie184* | *atl-1* (*ie148* [*atl-1 R1966K S1971A*]) V | *ieSi38*[*sun-1p::tir1::mRuby::sun-1 3'UTR*, *Cbr-unc-119(+)*] IV |

**Table S2.**
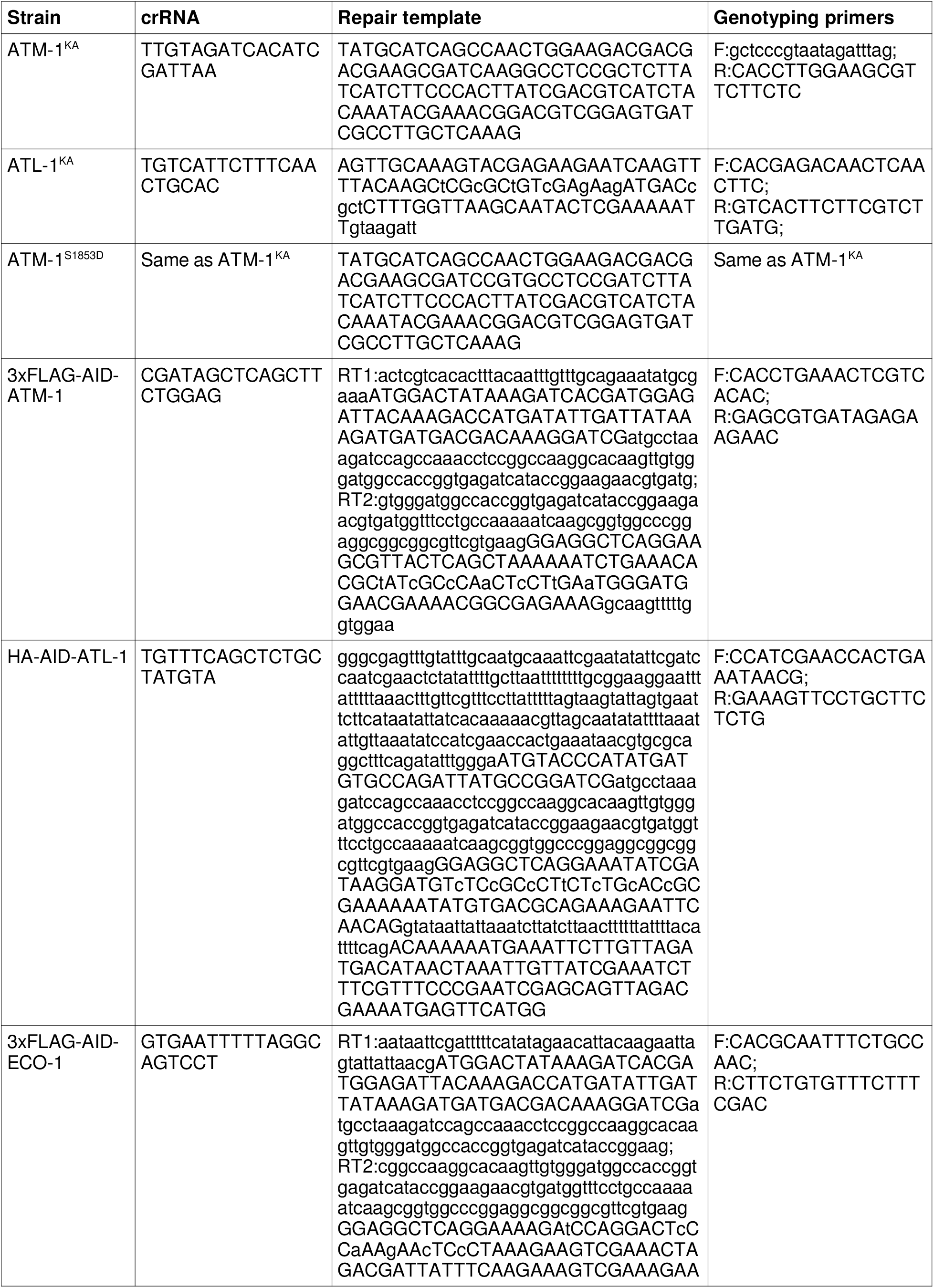
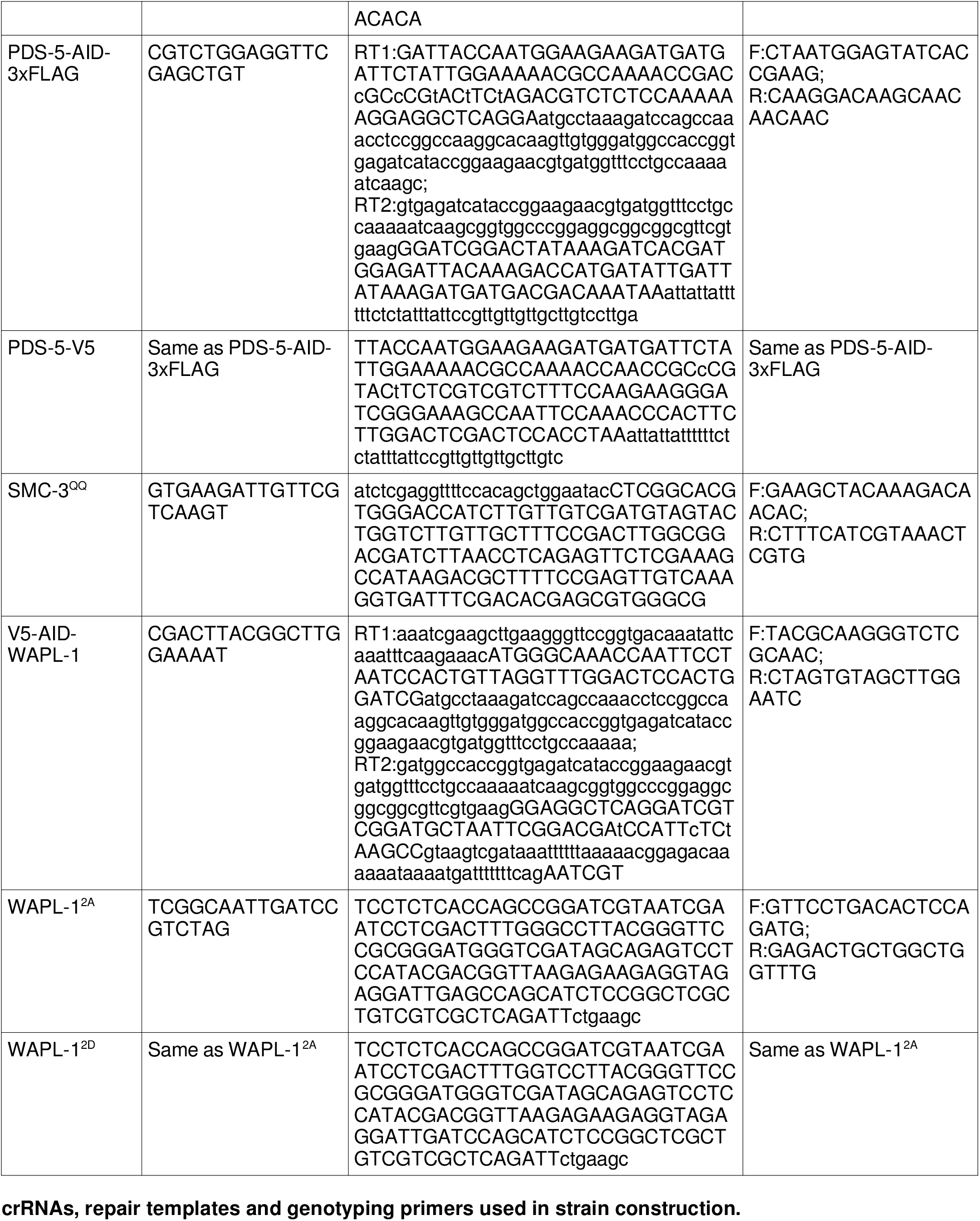
crRNAs, repair templates and genotyping primers used in strain construction.

**Table S3.**
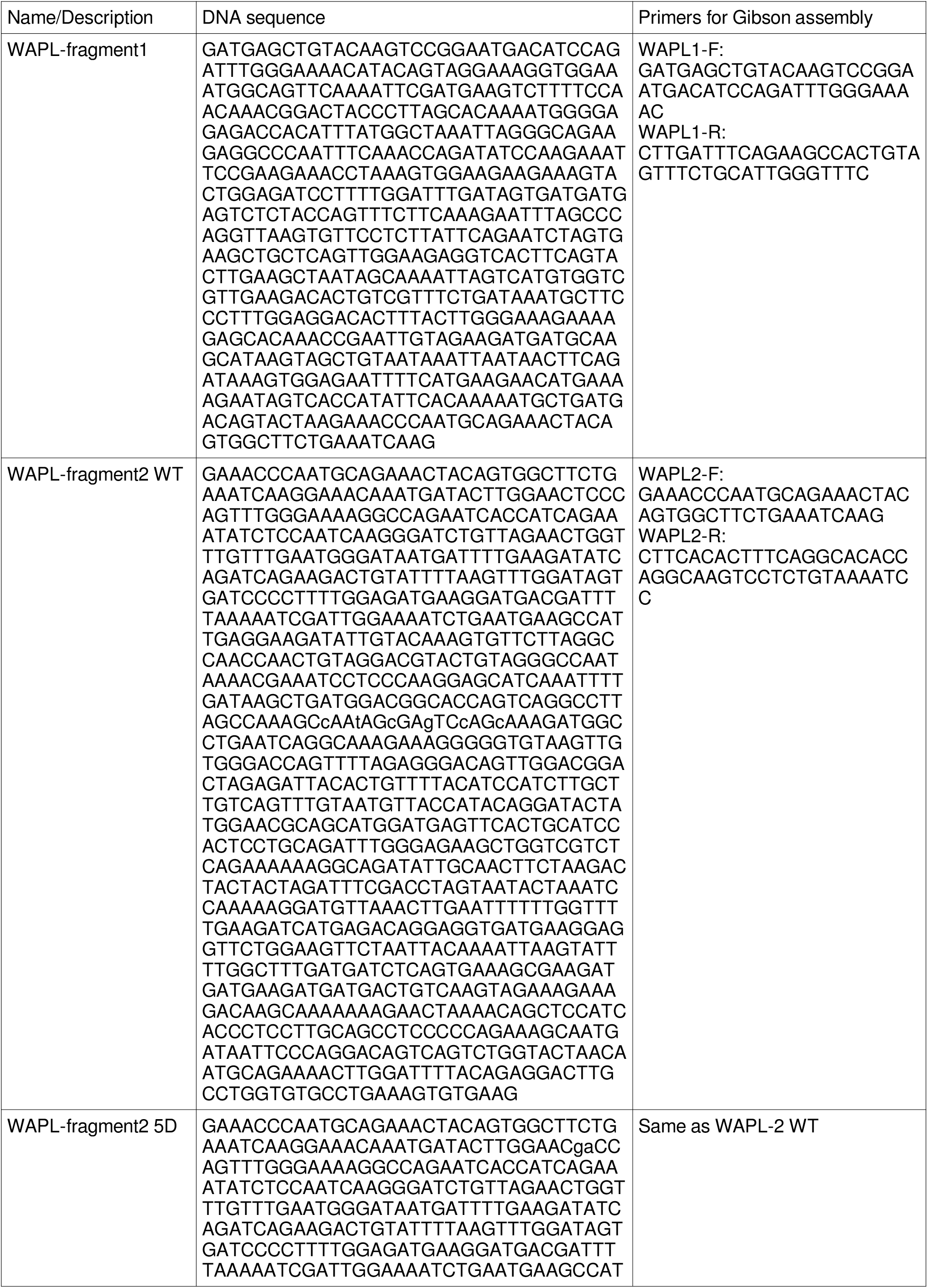
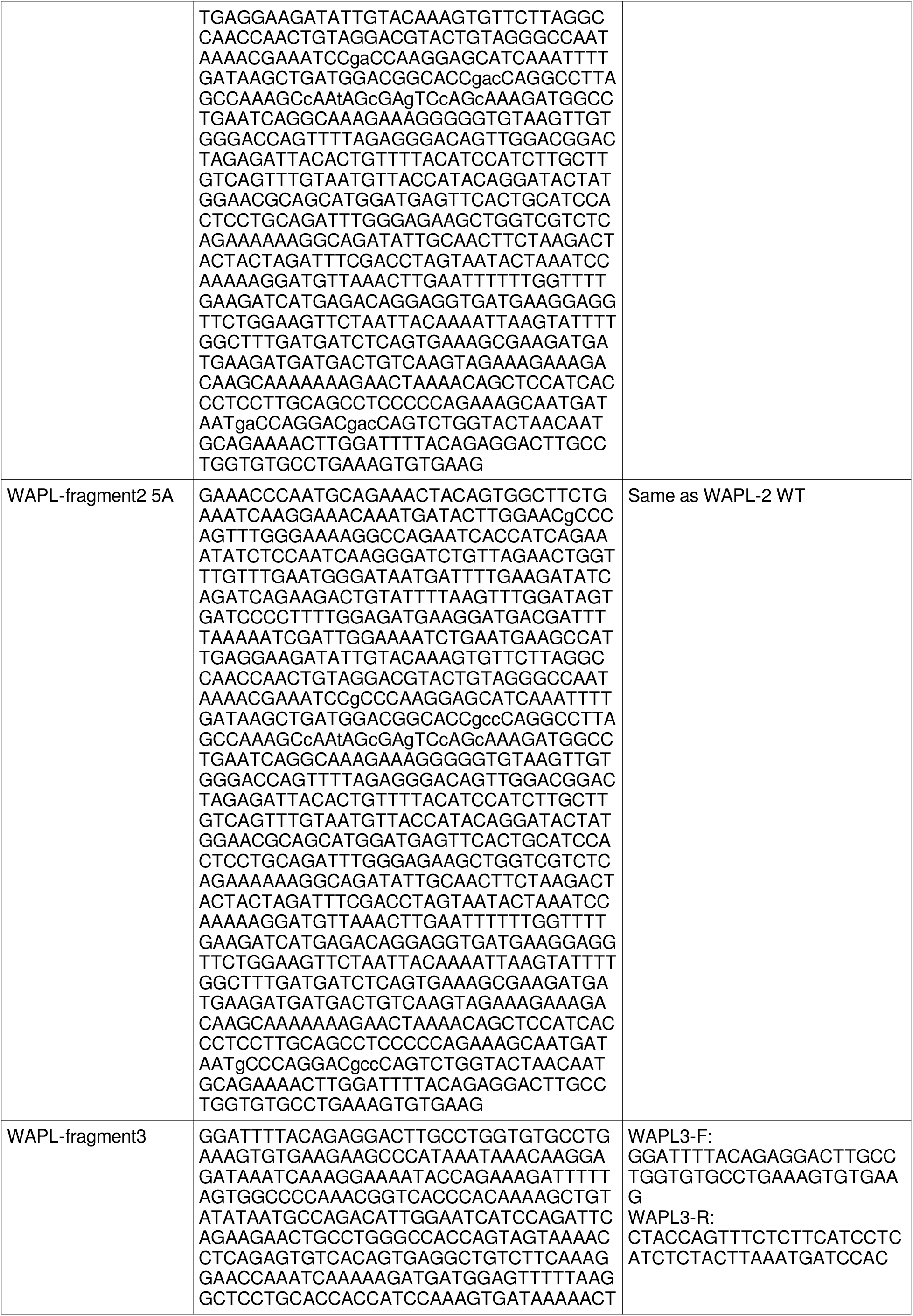
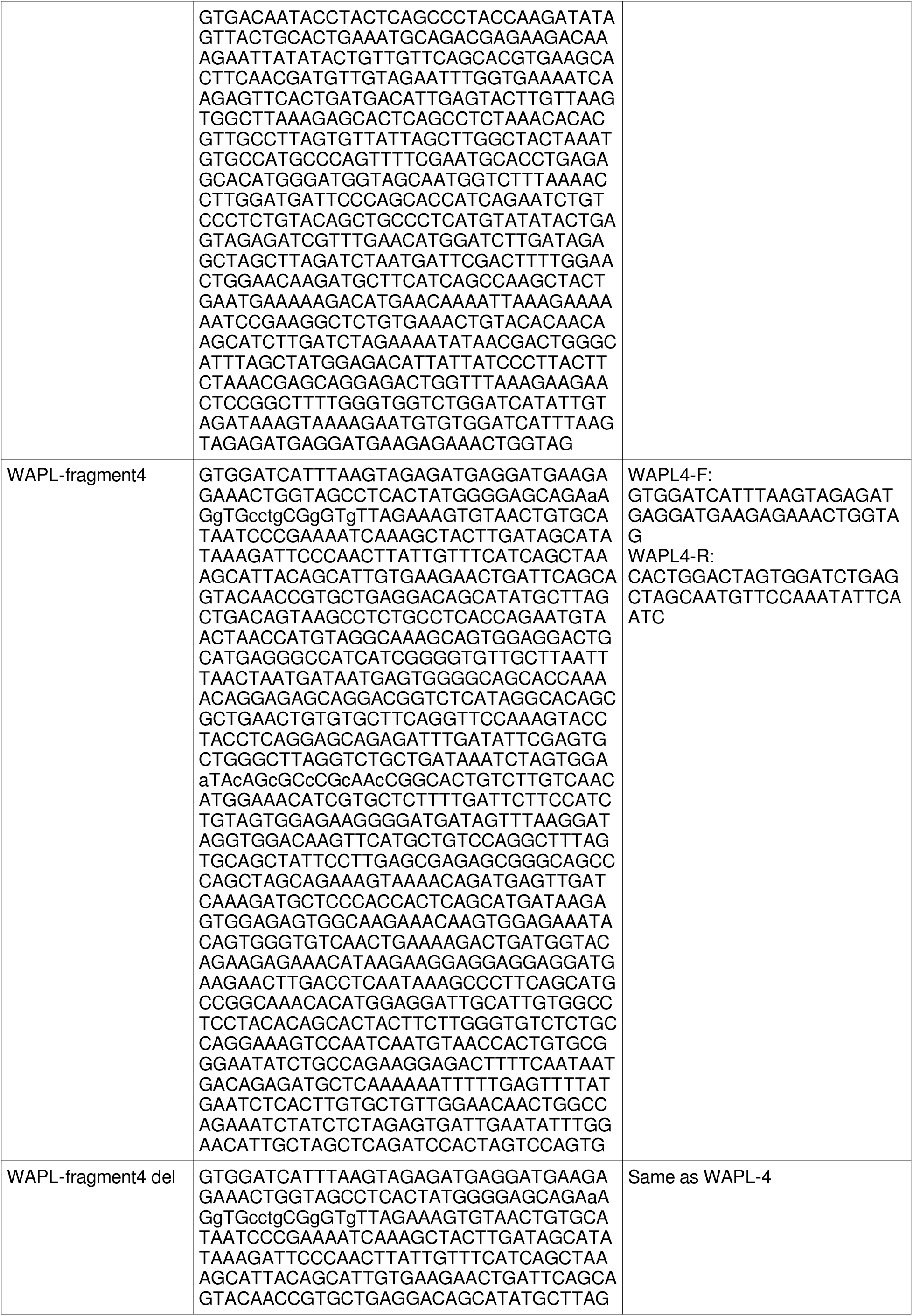
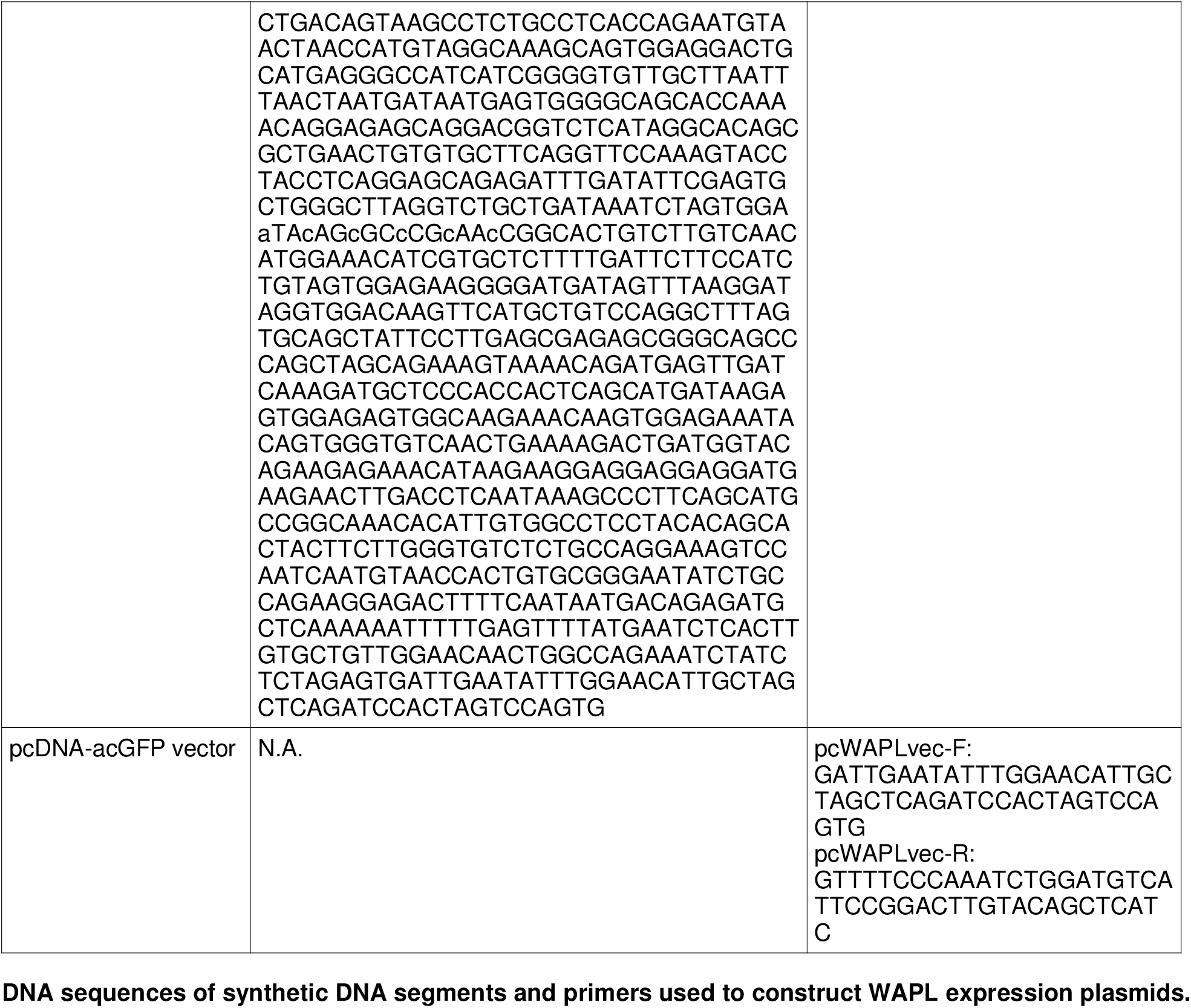
DNA sequences of synthetic DNA segments and primers used to construct WAPL expression plasmids.

